# Human *PRE-PIK3C2B* exhibits long-range intra- and inter-chromosomal interactions with genomic regions enriched in repressive marks

**DOI:** 10.1101/2020.11.11.378745

**Authors:** Jayant Maini, Ankit Kumar Pathak, Kausik Bhattacharyya, Narendra Kumar, Ankita Narang, Neha Jain, Inderpreet Singh, Vanika Dhingra, Vani Brahmachari

## Abstract

Human PRE-PIK3C2B is a dual nature polycomb response element that interacts with both polycomb as well as trithorax members. In the current study, using 4C-Seq (**C**apturing **C**ircular **C**hromosomal **C**onformation-**Seq**uencing), we identified long-range chromatin interactions associated with *PRE-PIK3C2B* and validated them with 3C-PCR. We identified both intra-as well as inter-chromosomal interactions, a large proportion of which were found to be closely distributed around transcriptional start sites (TSS). A significant number of interactions were also found to be associated with heterochromatic regions. Meta-analysis of ENCODE ChIP-Seq data identified an overall enrichment of YY1, CTCF as well as histone modification such as H3K4me3 and H3K27me marks in different cell lines. Almost 90% interactions were derived from either intronic or intergenic regions. among which large proportions of intronic interactors were either unique sequences or LINE/SINE derived. In case of intergenic interactions, majority of the interaction were associated with LINE/SINE repeats. We further found that genes proximal to the interactor sequences were co-expressed, they showed reduced expression. To the best of our knowledge this is one of the early demonstrations of long-range interaction of PRE sequences in human genome.

## 1.1 Introduction

Organization of the genome into higher-order chromatin architecture generates chromosome territories in the nucleus of a eukaryotic cell. This process involves extensive looping of chromatin fibers leading to the highly compacted organization [1]; [2]; [3]. This 3-dimensional organization not only solves the packing problem but also brings about co-regulation of physically distant genes and further, provides accessibility to the transcriptional machinery. The specific position of genes and chromosomes in the nucleus defines tissue- and cell type-specific gene expression [4]; [5]; [6]; [7]; [8]. Recently, with the development of Chromosome Conformation Capture assays (3C) and imaging techniques, a number of possibilities for the detection and study of the chromosome architecture has opened up [9]; [10]; [11]; [12]. 3C is an important technique that utilizes the concept of proximity ligation assay to capture genomic interactions [9].

A strong correlation is found between epigenetic modifications, chromatin compaction and transcription. It is well-known that most of the heterochromatic sites including the inactive X-chromosome in female mammals, are associated with the nuclear periphery and this, correlates with reduced expression of genes [13]; [14]; [15]; [16]. Furthermore, the territorial compaction is shown to be affected by epigenetic modifications such as histone acetylation [17]; [18].

A number of topologies resulting in the formation of DNA loops bring the target genes closer to the distal enhancers, silencers and insulators and facilitate various biological processes [19]. The 5’-3’ gene looping is important for transcription termination, while the PRE mediated looping is important for gene silencing, as it brings the polycomb response element closer to the promoter [19]. Thus, the identification of the cis-elements contributing to genome topology and long-range interactions are fundamental to understanding gene regulation holistically.

The Polycomb (PcG) complexes (PRC1 and PRC2) bind to the Polycomb Response Elements (PREs) and lead to the formation of H3K27me3, a repressive histone modification on the chromatin. While the Trithorax complexes are recruited to Trithorax Response Elements (TREs) and bring about H3K4me3, an activating histone modification. These histone marks are identified by various ATP-dependent chromatin remodelers that alter the structure of the chromatin to facilitate activation/ repression of gene expression. PRC1 complex is responsible for the compaction of nucleosomal arrays and exhibits both diffused and compact localization [20]. The PcG bodies are identified as nuclear domains enriched with heterochromatin regions that may be dispersed and not close to each other on the linear chromatin [21]. Boettiger and colleagues used the Oligopaint approach and classified the *Drosophila* genome into three major domains; the active, inactive and Polycomb-repressed domains where the chromatin is very densely packed [22]. These polycomb repressed regions were enriched at transcription factor coding genes involved in development [22].

The role of PREs in long-range chromatin interactions is well established in *Drosophila*, however, there are studies that establish insulator elements and not PREs as the major contributor to long-range interactions between polycomb targets [23]; [24]. Recent reports also provide evidence for cross-talk between strong and weak PREs in the regulation of gene expression through the maintenance of chromatin states [25]. Further, the role of polycomb members in long-range interactions is demonstrated in *Drosophila*, as in the case of Polyhomeotic (Ph) with mutation in the sterile alpha motif (SAM) which decreased the number of chromatin interactions disrupting the PcG clusters and in contrast, increase in Ph led to a significant increase in the number of clusters and chromatin interactions [26].

The long-range interactions in the human genome have been detected and analysed under the ENCODE project by 5C/Hi-C approach [27]; [28].

There are a limited number of reports on the long-range interactions with specific reference to PcG complexes and the PRE/TRE sequences and most of these are focused on *PRE/TREs* of *Drosophila*. Among the few PREs identified and characterized in the human genome, the human *PRE-PIK3C2B* is a dual function DNA-element interacting with the PcG complex as well as the members of the MLL complex. Human *PRE-PIK3C2B* undergoes changes in the histone modification marks following its interaction with Polycomb (PRC2) or MLL complex [29]; [30]. It is established that the binding of PRC2/MLL complex members is dependent on the relative concentration of each other; knock-down of PRC2 members, increases the recruitment of MLL1 on *PRE-PIK3C2B* and vice-versa [30]. Hence, *PRE-PIK3C2B* was one of the early reports of a dual function *PRE/TRE* element. The other example is the D4Z4 in FSHD, where shortening of the repeat leads to its activating effect [31]. In case of *hPRE-PIK3C2B*, we observe the coordinated expression of the genes *MDM1* and *PPP1R15Bthat* are away by 21.4kb and 11kb respectively from *PIK3C2B* and are on either side of *PIK3C2B* and are transcribed from opposite strands. We observed an increase in the expression of the three genes upon knock-down of *YY1* gene. Though, we could not rule out the presence of independent regulators, this suggested the possibility of *hPRE-PIK3C2B* controlling the three genes. There is a possibility that this effect might be the result of long-range interactions. In the light of these observations, we mapped the long-range interactions of human *PRE/TRE-PIK3C2B* using 4C-Sequencing technique.

## 1.2 Materials and Methods

### 1.2.1 Materials

DMEM High Glucose (Gibco™ 11995-065), DMEM low Glucose (Gibco™ 11885-084), opti-MEM 1 (11058-021) were obtained from Invitrogen. 0.5% Trypsin-EDTA (15400-054), Fetal Bovine Serum (Gibco™; 10270-106), Antibiotic-Antimycotic (Gibco 15240-062) and Nuclease free water were purchased from Invitrogen, USA. Lipofectamine 2000 (Gibco 116668-027) reagent was procured from Invitrogen. siRNA duplex for YY1

### 1.2.2 Methods

#### 1.2.2.1 Cell, transfection and RNA isolation and qPCR

The human cell line HEK203T, U87 and HeLa were propagated and harvested as in Maini et al., 2017. Transfection and RNA isolation, cDNA synthesis and qPCR were carried out as described in Maini et al., (2017) [30].

#### 1.2.2.2 Circular chromosome conformation capture assay (4C assay)

We followed the method described by Rawat et al., (2017) [32] which is a modification of the protocol described by van de Werken et al., (2012) [33]. The following sections described the protocol in brief.

#### 1.2.2.3 Cross linking of chromatin

The cells were resuspended in 1 X PBS and then divided into aliquots containing ~10 million cells in 15 mL falcon tubes. The volume was then made up to 10 mL. The cells were cross-linked with 1% formaldehyde (270 μL of 37% formaldehyde) for 10 mins at 25°C, followed by quenching of cross-linking reaction with 0.125 M for 10 min at 25°C. The suspension diluted with ice cold 1X PBS (volume made up to 20 mL) and centrifuged at 380 x g for 8 mins at 4°C.

#### 1.2.2.4 Preparation of nuclei

The pellet was resuspended in a 5mL ice-cold lysis buffer and homogenized with 15 strokes on ice. The cell suspension was kept at 4°C in an end-to-end rotor for 30 mins and centrifuged at 600 x g for 5 mins at 4°C. The pellet was washed with 1 mL of 1X restriction digestion buffer and the nuclear pellet was stored at −80°C.

#### 1.2.2.5 Restriction endonuclease treatment

The nuclei pellet was resuspended in 1.2X restriction enzyme buffer followed by incubation at 37°C on a shaker (1100 rpm) for 1 hour with 10% SDS incubated. The nuclei suspension was incubated with 20% Triton X100 for 1 hour at 37°C on the shaker. 100 μL of sample was aliquoted as the undigested control. The rest was incubated with 800U of restriction enzyme (Csp6I) overnight at 37°C on the shaker (1100 rpm). Following this, 100 μL of sample was aliquoted as a digestion control. The DNA was isolated from both undigested and digested control by Phenol-Chloroform-Isoamyl alcohol (PCI) extraction and ethanol precipitation. The digested sample was analyzed by electrophoresis on 0.8% agarose gel for digestion efficiency (Figure 1A).

**Figure 1.**
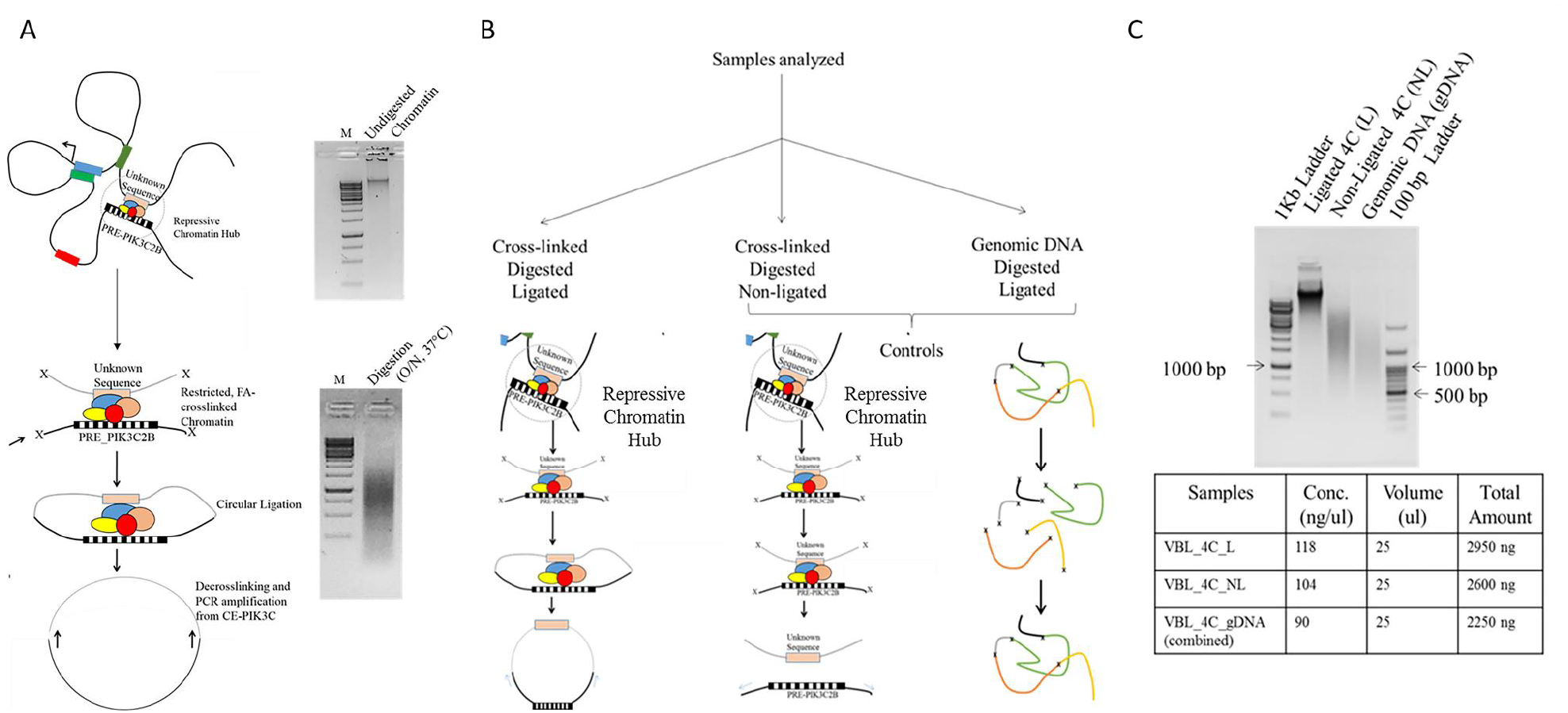
Schematic of 4C assay. A. Formaldehyde based cross-linking of cells followed by enzyme digestion and intra- and inter-molecular ligation. Primers mapping to the bait sequence were used to amplify the unknown sequences (Query sequences). B. The controls used to rule out false positive interactions. C. Electrophoresis profile showing the quality of 4C-samples used for sequencing. The table below shows the concentration and the total amount of samples used for 4C sequencing.

#### 1.2.2.6 Ligation

The nuclei lysate was diluted to 1ng/μL followed by addition of 10X T4 DNA ligase buffer (final concentration of 1X), 10% triton X100 (final concentration of 1%), 10 mg/mL BSA (final concentration of 0.1mg/mL) and 3000 units of T4 DNA ligase. The volume was made up to 8 mL. The ligation reaction was incubated at 4°C for 3 days with continued renewal of ATP every second day. The samples were then treated with proteinase K (50 uL of 10 mg/mL) overnight at 65°C. The samples were purified using PCI extraction and ethanol precipitation. The pellet was resuspended in 100 ul of Nuclease-free water and the purified ligated DNA was treated with 1ul of 10 mg/mL RNase A for 15 mins at 37°C. The ligated sample was stored in −20°C. The library was checked on 0.8% agarose gel (Figure 1C). For each set of reaction, two controls were set up. One was the digested, non-ligated control and the other genomic DNA (gDNA) digested and ligated (to get rid of non-specific interactions) (Figure 1B and C).

#### 1.2.2.7 PCR amplification and 4C sequencing

200 ng of 4C DNA was used as the template in each PCR. The PCR reaction and conditions has been mentioned in **Supplementary Table 1–4**

#### 1.2.2.8 4C sequencing

The amplicons were then sequenced using the Illumina (MiSeq and NextSeq platform). Three set of samples were subjected to 4C analysis as follows: (i) L-the cross linked, CspI digested and ligated, (ii) NL-cross-linked, CspI digested but non-ligated (control 1) and (iii) gDNA-HEK293T genomic DNA which was CspI digested and then religated (control 2). The controls were used to exclude false positive interactions. Sequencing adapter contamination was removed from the sequencing data using Trimmomatic (0.36) software (Bolger et al., 2014). Quality checks of sequencing reads were performed using FastQC (https://www.bioinformatics.babraham.ac.uk/projects/fastqc/). The reads were aligned to the human genome (hg38) using Bowtie2 [34]. The number of reads obtained from sequencing experiments on Illumina MiSeq and NextSeq for different samples is listed in **Supplementary Table 5** and **6**

#### 1.2.2.9 4C Sequence data clean-up and analysis

The sequencing reads from both the platforms were combined for futher analysis (**Supplementary Table 7**). Following this the primer sequences (upto restriction site) were trimmed from 5’ ends and 3’ end of reads (viewpoint sequence). The above two steps ensure only sequences from the interacted regions are retained for alignment. Sequencing adapter contamination was removed using trim_galore (https://www.bioinformatics.babraham.ac.uk/projects/trim_galore/). The number of reads after trimming the adaptor sequence in different samples ranged from 2.8-4.4X10^5^ (Table 1). The reads for each experiment were aligned to hg19 using bowtie2 (Table 2). Detection of DNA interacting regions with viewpoint was performed using the r3Cseq package [35] using default setting. A summary of the number of detected interacting regions is as shown in Table 3. Q-value is for False Discovery Rate [35].

**Table 1.** Summary of remaining sequences after trimming

| Files | Number of Reads |
| --- | --- |
| SO_5382_VBL_4C_gDNA_R1_val_1.fq | 286703 |
| SO_5382_VBL_4C_gDNA_R2_val_2.fq | 286703 |
| SO_5382_VBL_4C_L_R1_val_1.fq | 302991 |
| SO_5382_VBL_4C_L_R2_val_2.fq | 302991 |
| SO_5382_VBL_4C_NL_R1_val_1.fq | 440105 |
| SO_5382_VBL_4C_NL_R2_val_2.fq | 440105 |

**Table 2.**
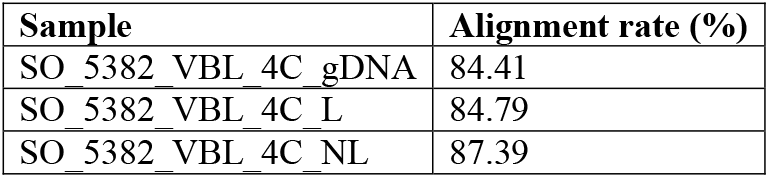
Alignment summary

| Sample | Alignment rate (%) |
| --- | --- |
| SO_5382_VBL_4C_gDNA | 84.41 |
| SO_5382_VBL_4C_L | 84.79 |
| SO_5382_VBL_4C_NL | 87.39 |

**Table 3.**
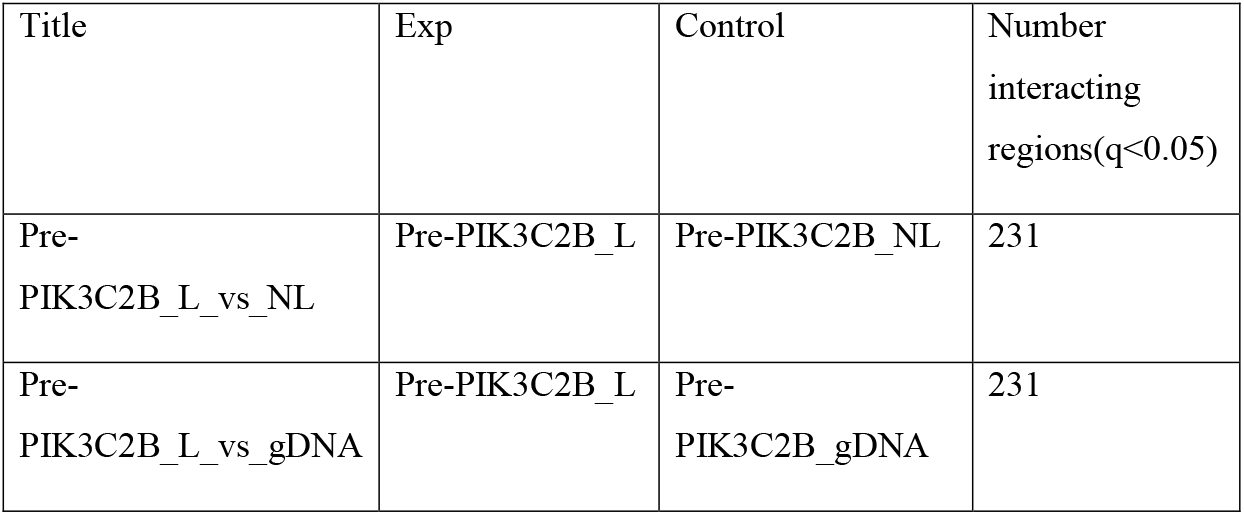
Number of interacting regions identified

We also checked the accuracy of the protocol we used, by performing 4C using the same protocol but with BamH1 restriction enzymes rather than Csp6I, to map the already known interaction at the *H19-IGF1* locus (Supplemental Figure S1).

## 1.3 Results

### 1.3.1 Identification of interactor query sequences

After aligning the paired reads to the human genome, we selected the genomic regions for further analysis based on two criteria: (i) the coverage ≥5X and (ii) their representation only in the ligated (L) sample and not in non-ligated (NL) or digested and re-ligated gDNA. The common sequences between the L, NL and gDNA were considered as false positives and were not analysed further. They can also be the result of re-ligation because of proximity to the *hPRE-PIK3C2B*, the bait sequence. Additionally, only the regions with ≥500 reads (≥ 100 reads per million), were taken for further analysis.

### 1.3.2 Identification of intra-and inter-chromosomal interactions

We detected 231 interactors, of which 22 were intra-chromosomal and 209 interactors were inter-chromosomal (Figure 2A). The query sequences varied in length from 60 to 200 bp (Figure 2B). The intra-chromosomal sequences that interact with the bait *PRE-PIK3C2B*, map to regions that are approximately 30-150 Mb from the bait sequence (Figure 3A).

**Figure2.**
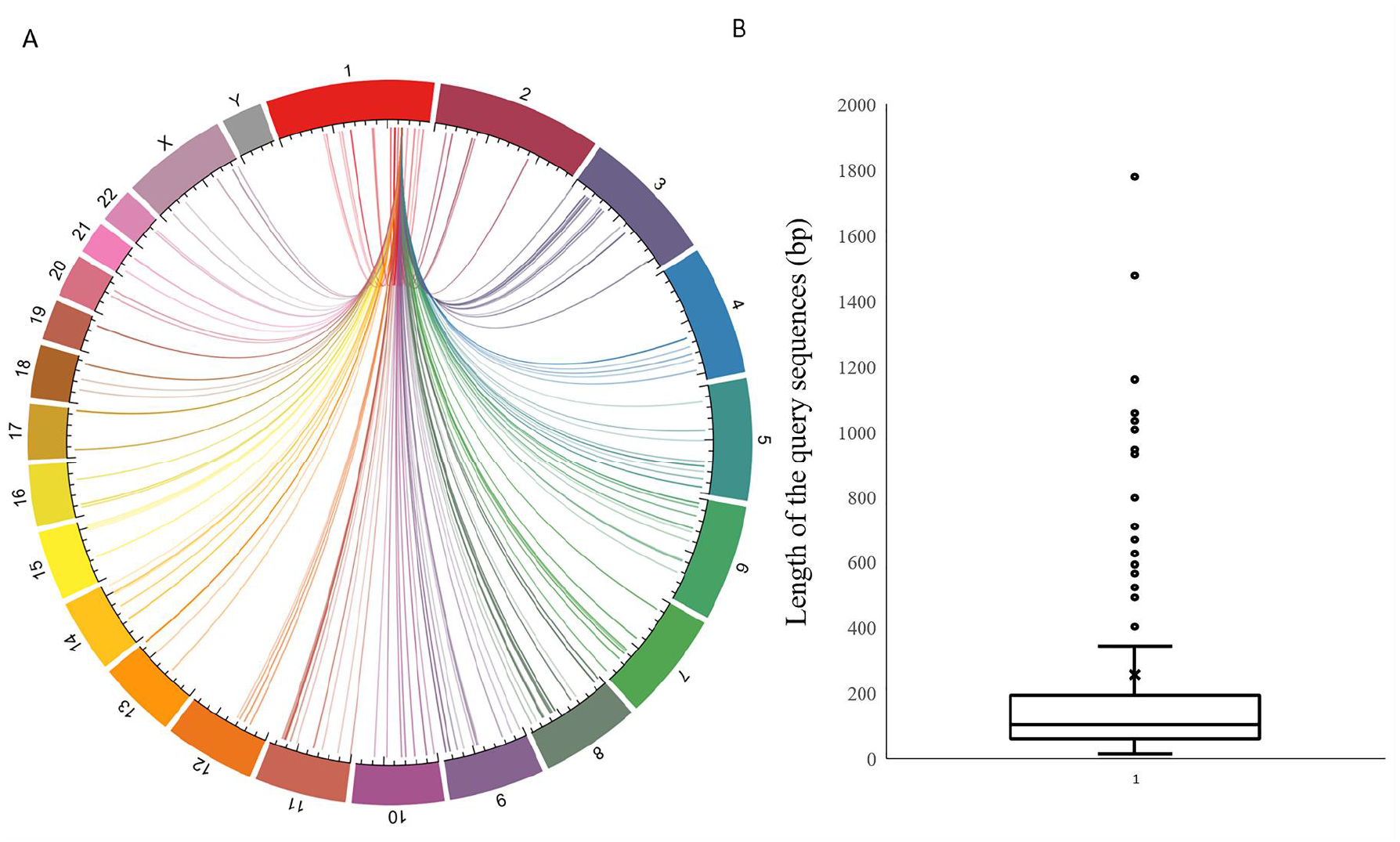
Mapping of the interactor sequences on human genome. A-the circos plot indicating chromosome-wise distribution of the interactor sequences. The chromosomes are marked 1 to 22, along with the X and Y, B - Distribution of the interactors based on their length.

**Figure 3.**
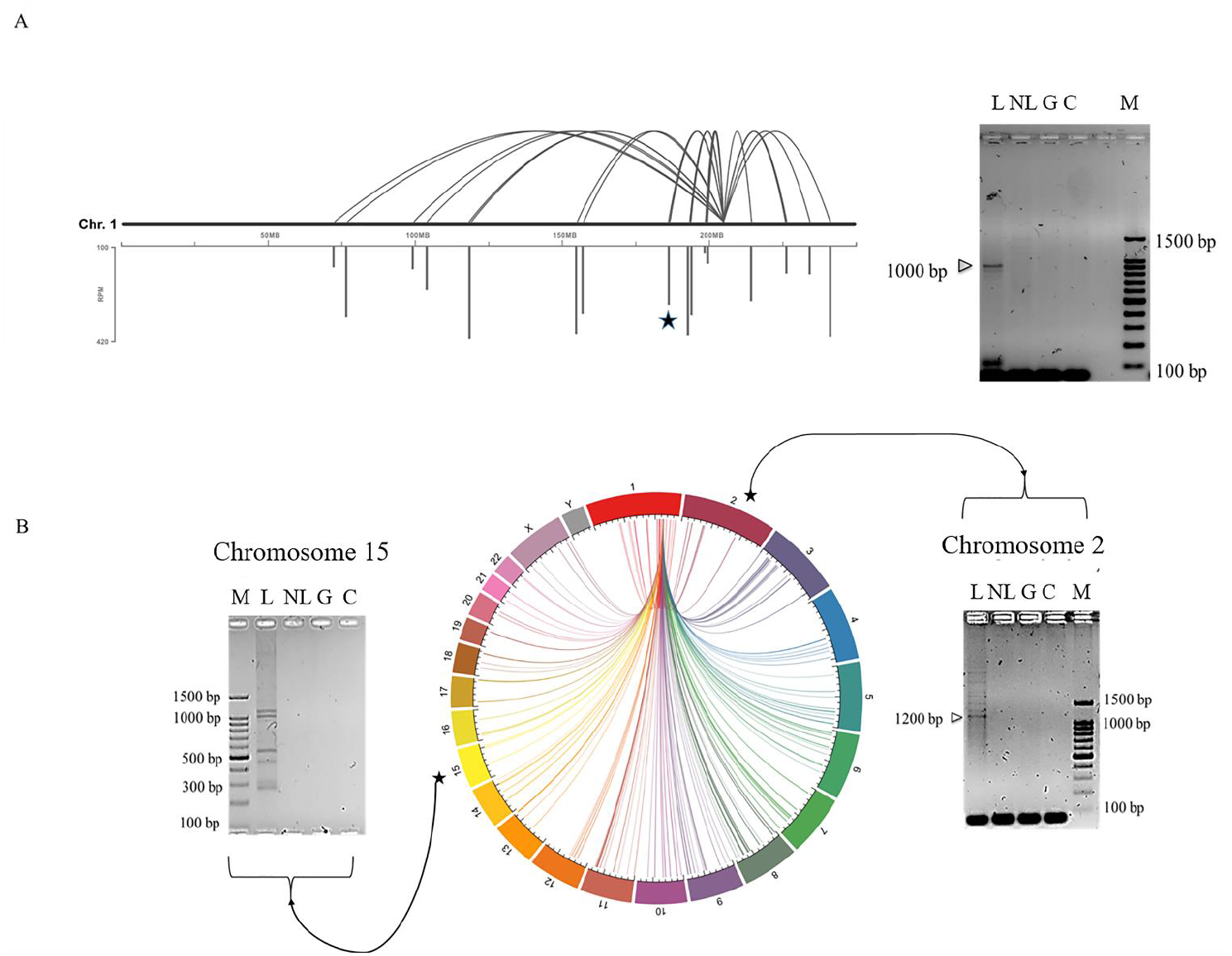
Validation of interactions by 3C followed by PCR. A-Intra-chromosomal (Chr1) interactions are indicated and the interactor, B-Indicates the position of the inter-chromosomal interactions (chr2 and chr15) chosen for validation. Interactors validated are marked with a star. The amplicons obtained are shown in the corresponding electrophoresis profile; L-ligated, NL-non-ligated, gDNA-genomic DNA restriction digested and ligated, C control lacking the template, M-100bp DNA ladder.

### 1.3.3 Validation of the intra- and inter-chromosomal interactions using 3C approach

The validation of the long-range interactions was performed using a 3C approach whereby primers mapping to both the *hPRE-PIK3C2B* (bait) and the interacting sequences identified by 4C, were used to amplify the interacting regions. Using 3C, 6 interactions were validated. A representative figure of intra-chromosomal and inter-chromosomal interactions is shown in Figure 3. These interactions were found to be specific for the ligated sample i.e. their amplicons were not detected in non-ligated and gDNA (digested and re-ligated) controls. The validated intra-chromosomal interaction was at a distance of approximately 40 Mb from the *PRE-PIK3C2B* (bait) sequence (Figure 3A). The inter-chromosomal interactor regions identified, mapped to different chromosomes such as 2, 4, 13, 15 and X. Upon validation of interactions, multiple amplicons were observed on chromosomes 4, 13, 15 and X, indicating that these inter-chromosomal interactor sequences might form a part of many interactions with multiple interacting partners (Figure 3B and Supplementary Figure S2 & S3).

### 1.3.4 Distribution of interactor sequence around TSS

The interactor sequences identified were found to be distributed around the Transcription Start Sites (TSS), with almost 193 interactions (out of the total 231) mapping to regions within +/- 2Mb of the TSS. Upon detailed analysis, it was identified that 113 interactions within the +/- 2Mb mapped to the region with +/- 50Kb of the TSS, which account for almost 58% of the interactions mapping within +/- 2Mb (Figure 4). This shows that a number of interacting partners are present in close proximity to the TSSs thereby pointing towards their potential role in coordinated gene expression

**Figure 4.**
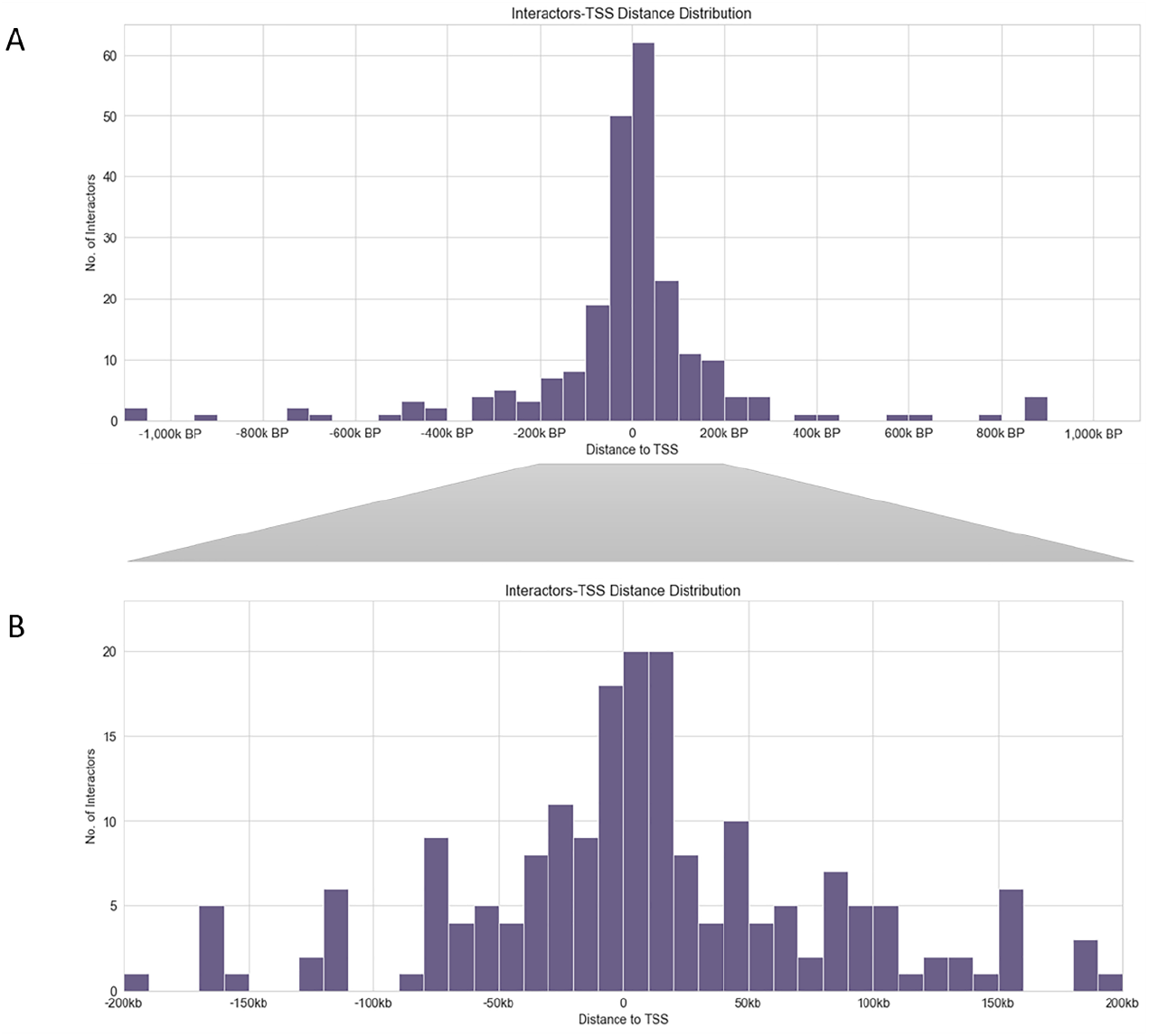
Distribution of interactors around the TSS. A-distribution of interactor sequences around TSS in +/- 1000Kb window, B- the same in +/- 200 kb window. Please see methods section for details.

### 1.3.5 Expression analysis of genes associated with Interactor sequences

A number of interactor sequences mapped within the genic regions or mapped to intergenic regions that were in close proximity to transcription start sites of the genes. Since these sequences interact with *PRE-PIK3C2B*, the genes associated with interactor sequences could be co-regulated and thus, co-expressed. In order to investigate this, the genes were selected on the basis of their proximity to most significant interactor regions followed by comparative expression analysis. The expression data in the form of TPM (Transcripts per Million) was downloaded from the online gene expression database, GEPIA (Gene Expression Profiling Interactive Analysis) [36] and compared (Figure 5A & B). The TPM value for *PIK3C2B* was also included and *GAPDH* was used as a control. In addition, the expression of a random set of genes, not associated with any of the interactor sequences, was also selected for comparison (Figure 5B). This analysis clearly indicates that the genes associated with interactor sequences were co-expressed as compared to the uncoordinated expression of the random set of genes (the experimental control) (Figure 5A & B). The random set was an unbiased selection of genes that were not found to be associated with the interactor regions. The experimental gene set was found to be co-expressed in all the tissues selected for analysis (Figure 5).

**Figure 5.**
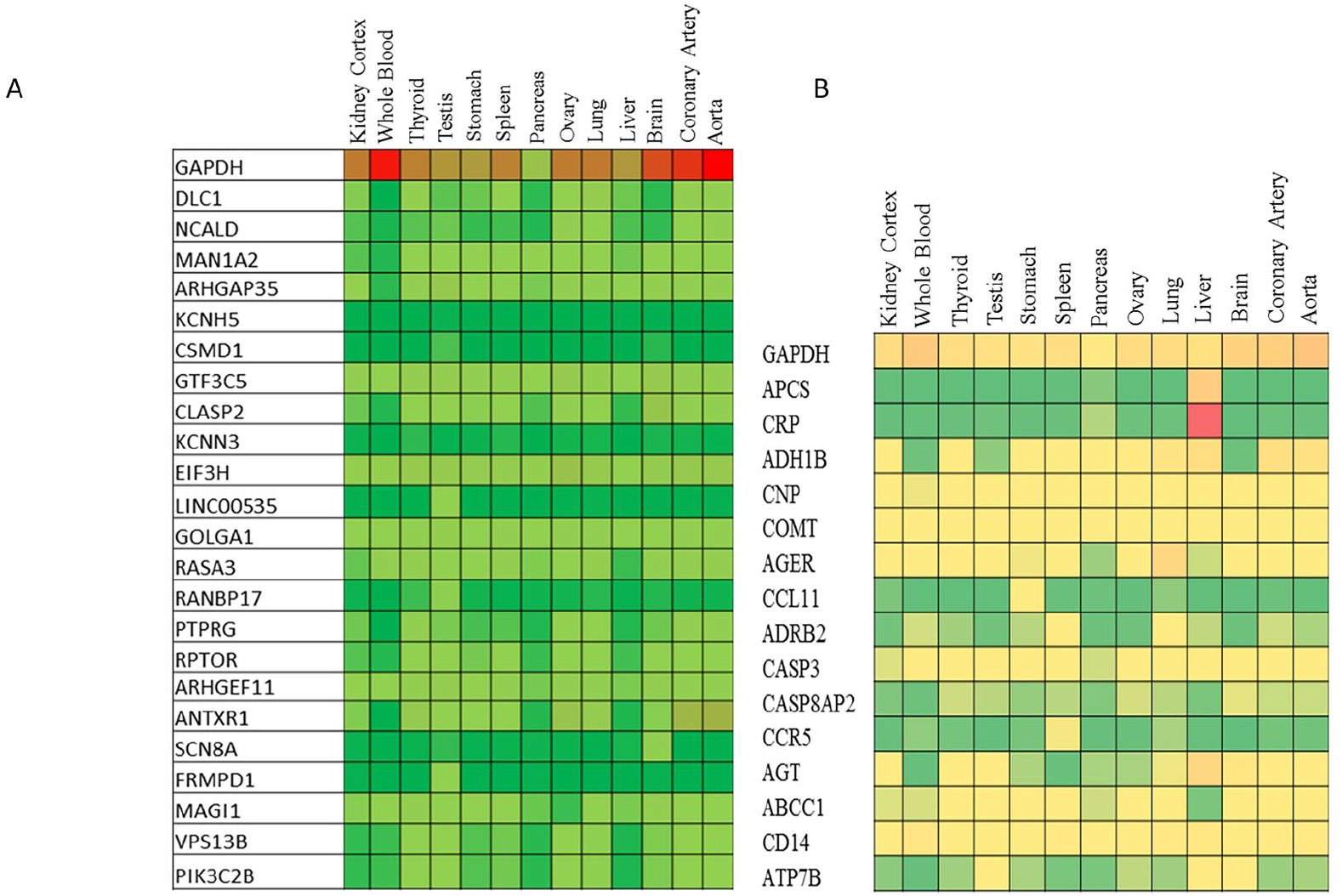
Heat map for expression of genes present in close proximity to the interactor sequences. Expression in terms of Transcripts per million (TPM). A-Gene expression analysis of interactor sequence-associated genes, B-Gene expression analysis of randomly selected genes, GAPDH is used as a housekeeping control in both A and B. The data was retrieved from GEPIA (Tang et al., 2017).

Using GEPIA, a gene expression correlation analysis was performed between a selective set of genes and *PIK3C2B*. These genes were shortlisted from the genes represented in the heatmap in Figure 5. This gene correlation analysis takes into account all the cell and tissue types. As shown in Figure 6, *DLC1, MAN1A2* and *ARHGEF11* show a very strong correlation in terms of expression with *PIK3C2B*. On the other hand, low positive correlation exists between *PIK3C2B* and *NCALD* & *ARHGAP35* (Figure 6).

**Figure 6.**
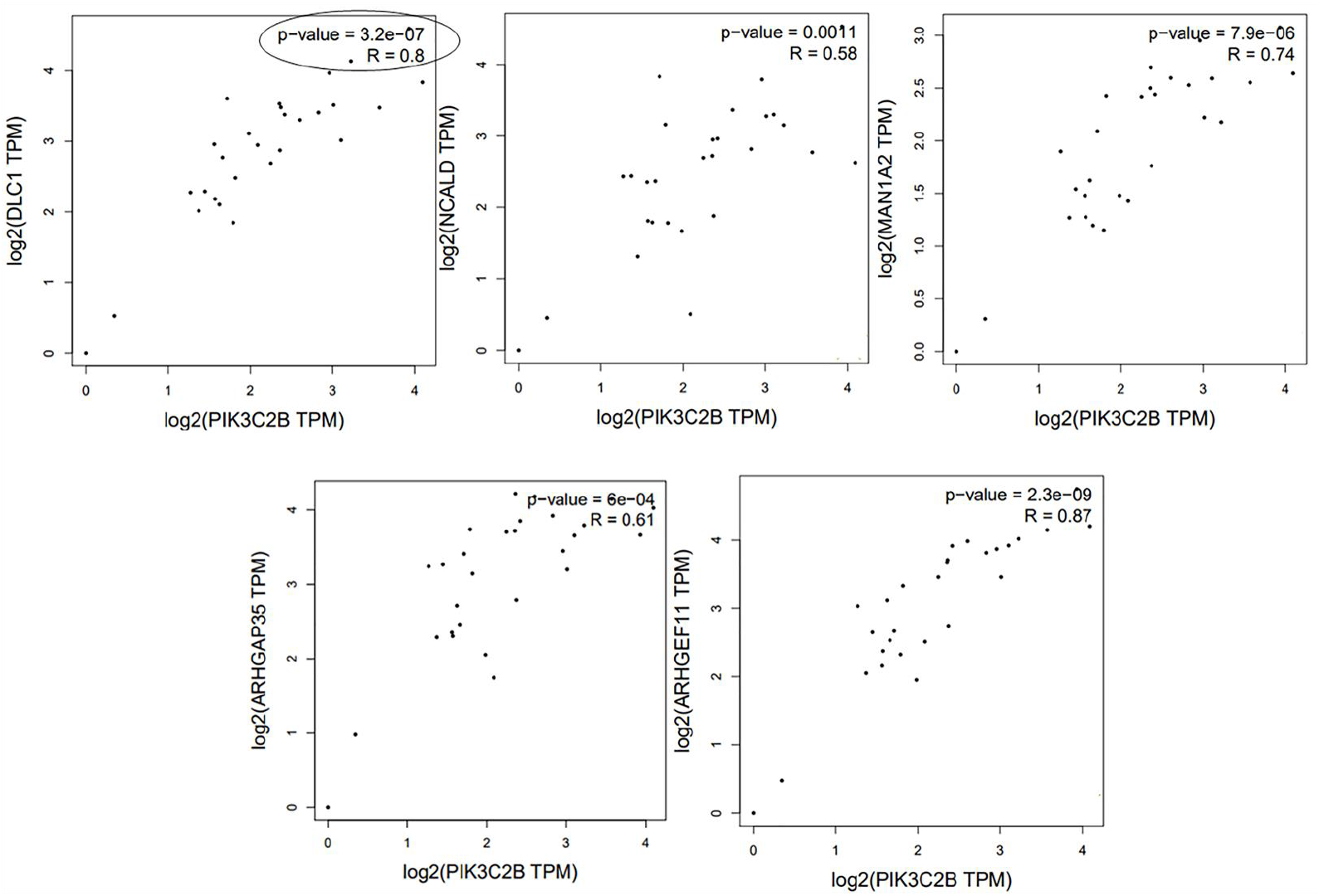
Analysis of correlation of expression. The expression of genes associated with interactor sequences is compared with that of PIK3C2B within which *PRE-PIK3C2B*is mapped. The expression data was retrieved from GEPIA (Tang et al., 2017).

### 1.3.6 Chromatin states associated with interactor sequences

Since very limited data is available for HEK cells in ENCODE, the query sequences were therefore mapped to the ENCODE ChIP-seq data for K562, HepG2, H1_hESC and GM12878 cell lines, to examine the chromatin state of the interactors (Figure 7A and B). In all the 4 cell lines, interactor sequences were predominantly associated with heterochromatin; out of (total number of interactions - 231), 140 interactor regions in K562, 150 in HepG2 and GM12878 and 160 in H1-hESCs (Figure 7A). A considerable proportion (5-40 interactions i.e. approximately 2-17% of total interactors) also mapped to various regions with other chromatin states such as active, weak or poised promoter, strong or weak enhancers, insulators and strongly or weakly transcribed regions. A small proportion (2-6%) of the interactors were found to be associated with Polycomb-repressed chromatin state (Figure 7B). Despite the different origin of the 4 cell lines, there is no significant difference in the number of interactors mapping to various chromatin states in these cell lines except the chromatin states associated with repression, weak transcription and transcription elongation (Figure 7B). Only GM12878 (lymphoblastoid cell line) and K562 (myelogenous leukemia cell line) are found to be related. However, the chromatin state of the interactors might vary depending on the cell line and its transcriptional profile.

**Figure 7.**
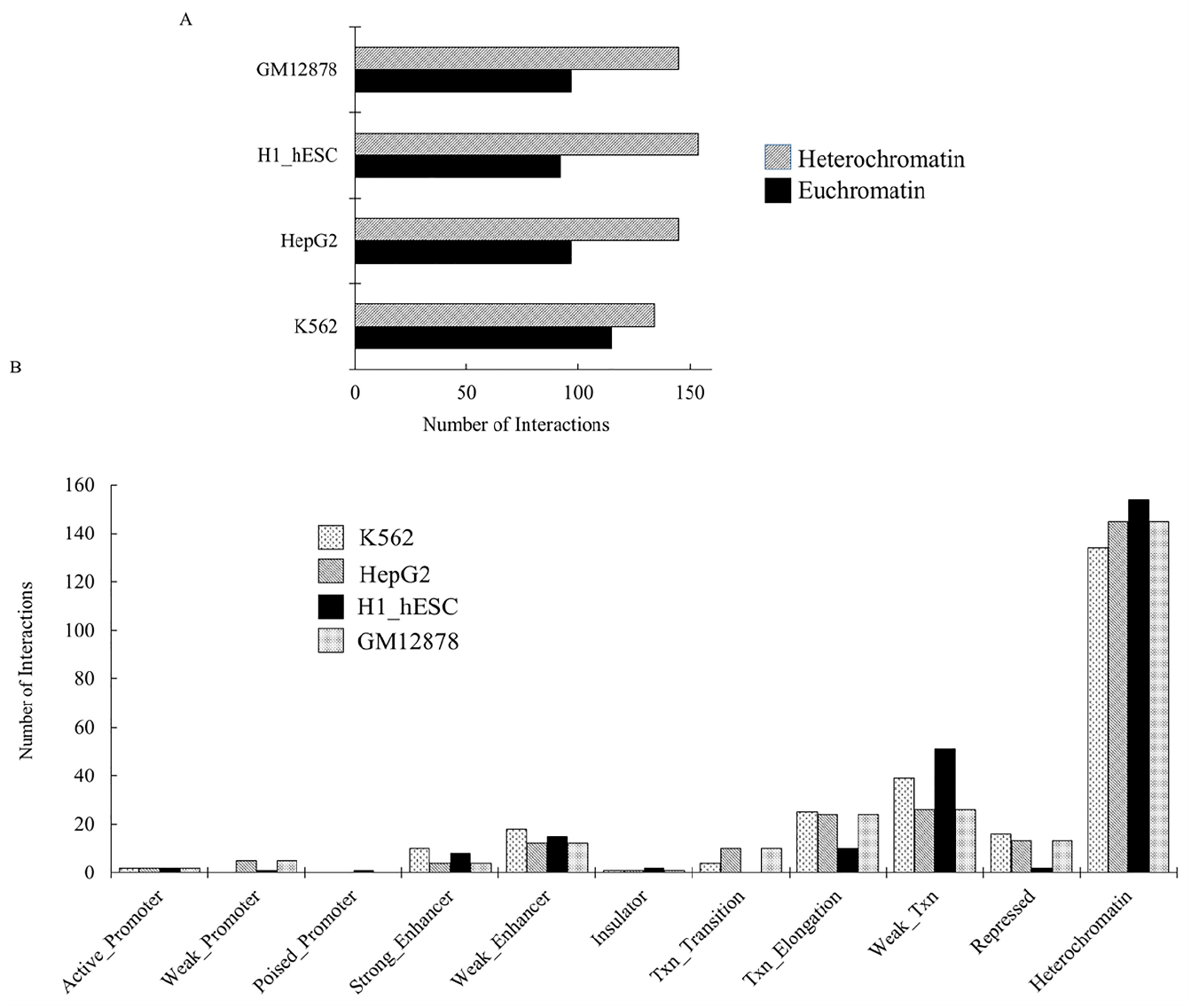
Chromatin state of the interactor sequences. A. Comparison of the chromatin state of interactor sequences in four different cell lines, B. Distribution of interactor sequences into different chromatin states in four different cell lines. The terms used to describe the chromatin states are as given by ENCODE (Encode Project Consortium, 2012).

### 1.3.7 Histone profile of interactor sequences

The histone profile maps were generated for interactor sequences as well as flanking 10kb upstream and 10kb downstream regions, using the ENCODE ChIP-seq data for various histone modifications (H3K4me3, H3K9me3, H3K27me3 and H3K36me3) and the epigenetic modifiers (EZH2, SUZ12, CBX2, YY1, CBX3 and CBX8) for the cell lines HeLa-S3, HepG2 and K562. The data for HEK293 and MCF-7 was limited and hence not analyzed. In the HepG2 cell line, there is an overall enrichment of H3K4me3 around the interactor sequences as observed in the +/- 2kb window, although other histone modifications are also present (Figure 8A). In contrast, the histone map for flanking regions in the K562 cell line indicates an overall enrichment of H3K27me3 especially within +/- 2kb window, although the H3K4me3 is also found to be associated with this region (Figure 8B).

**Figure 8.**
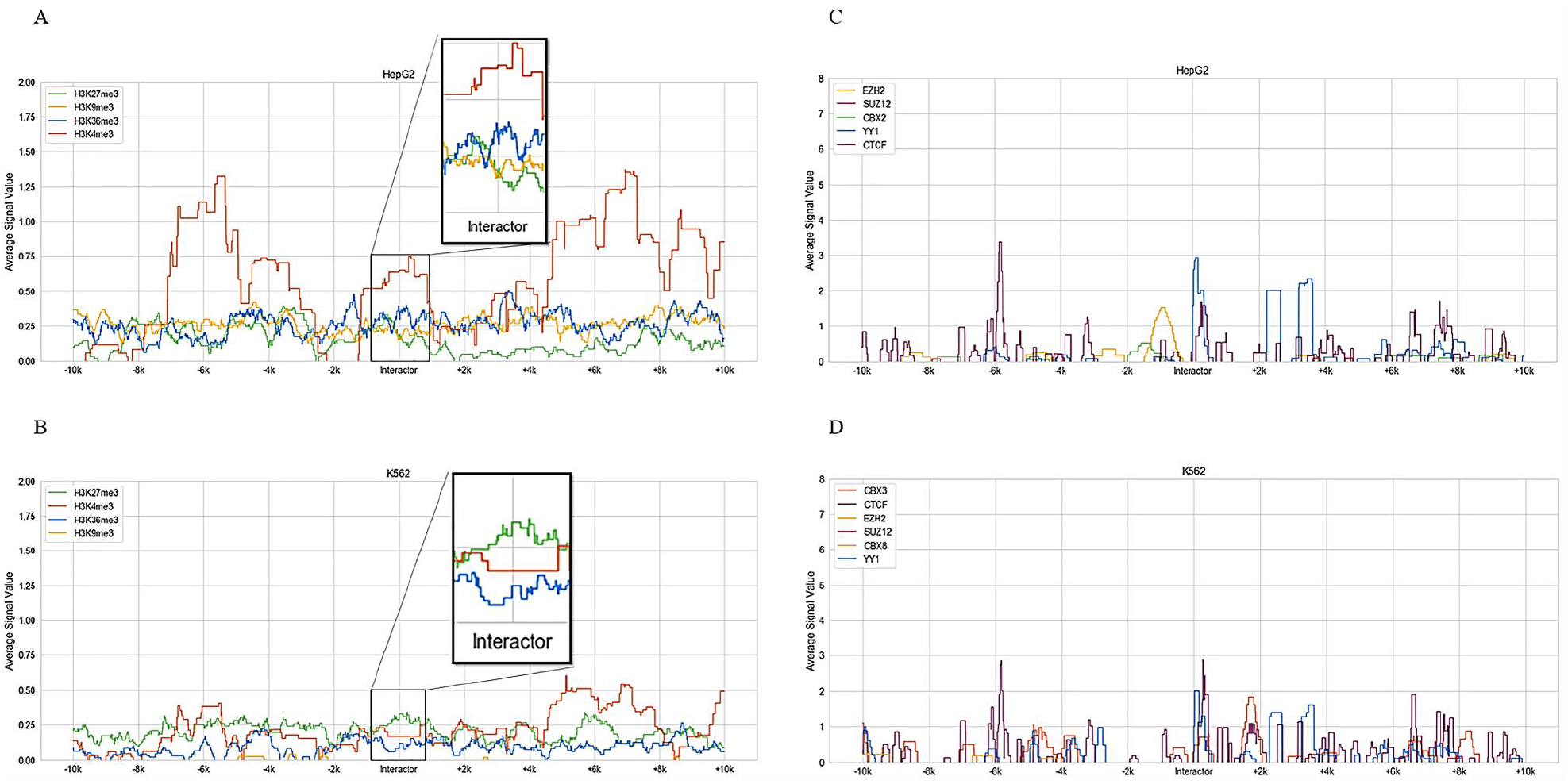
Profile of histone modification and epigenetic factor enrichment in regions flanking the interactor sequences. The data from K562 and HepG2 cell lines were taken from ENCODE. A and B-modified histone profile for regions flanking (±10Kb window) the interactor sequences, C and D -Profile of enrichment of epigenetic factors for regions flanking the interactor sequences (±10Kb window). The cell lines, modified histones and epigenetic factors analysed are given in each frame.

Epigenetic modifier-enrichment profile maps were generated for HepG2 and K562 cell lines (Figure 8C and D). In HepG2, the regions downstream to the interactor sequence (approximately +500 bp), were enriched in YY1 and CTCF binding, the enrichment of YY1 being greater than that of CTCF (Figure 8C). In K562 cells, there is an overall enrichment of YY1, CTCF and CBX3, 500bp downstream of the interactor sequence and the enrichment of CTCF is greater than that of YY1 (Figure 8D). The enrichment profile for the interactor sequences for HepG2 and K562 indicate an enrichment of YY1 binding, suggesting a repressive state.

K562 cell line shows a strong correlation between histone modification and epigenetic modifier enrichment, while HepG2 cell lines show increased enrichment of H3K4me3, as active histone mark, along with YY1 and CTCF enrichment, correlated with transcription repression and chromatin compaction.

### 1.3.8 Repetitive elements associated with interactor sequences

50% of the total interactions were intergenic and the rest of them mapped to the genic regions. These were further classified into intronic, exonic, 3’UTR and transcription termination sites (TTS) (Figure 9). The intergenic as well as genic interactors were further classified on the basis of their association with the repeat sequences (Figure 9). A majority of the intronic interactors were unique sequences (59 out of 108), and approximately 45% of the interactors (49/108) mapped to repetitive regions derived from LINE, SINE, LTR or simple repeats like (CA)n (Figure 9). Similarly, 40% of the intergenic interactors (46 out of 114) were found to be associated with unique sequences. The proportion of intergenic interactor sequences mapping to the repetitive regions (derived from LINE. SINE, LTR and AT-rich/low complexity repeats) is high at 60% (68 out of 114; Figure 9). Thus, the intergenic interactor sequences were enriched with repetitive sequences whereas the majority of the intronic interactors were unique sequences (Figure 9).

**Figure 9.**
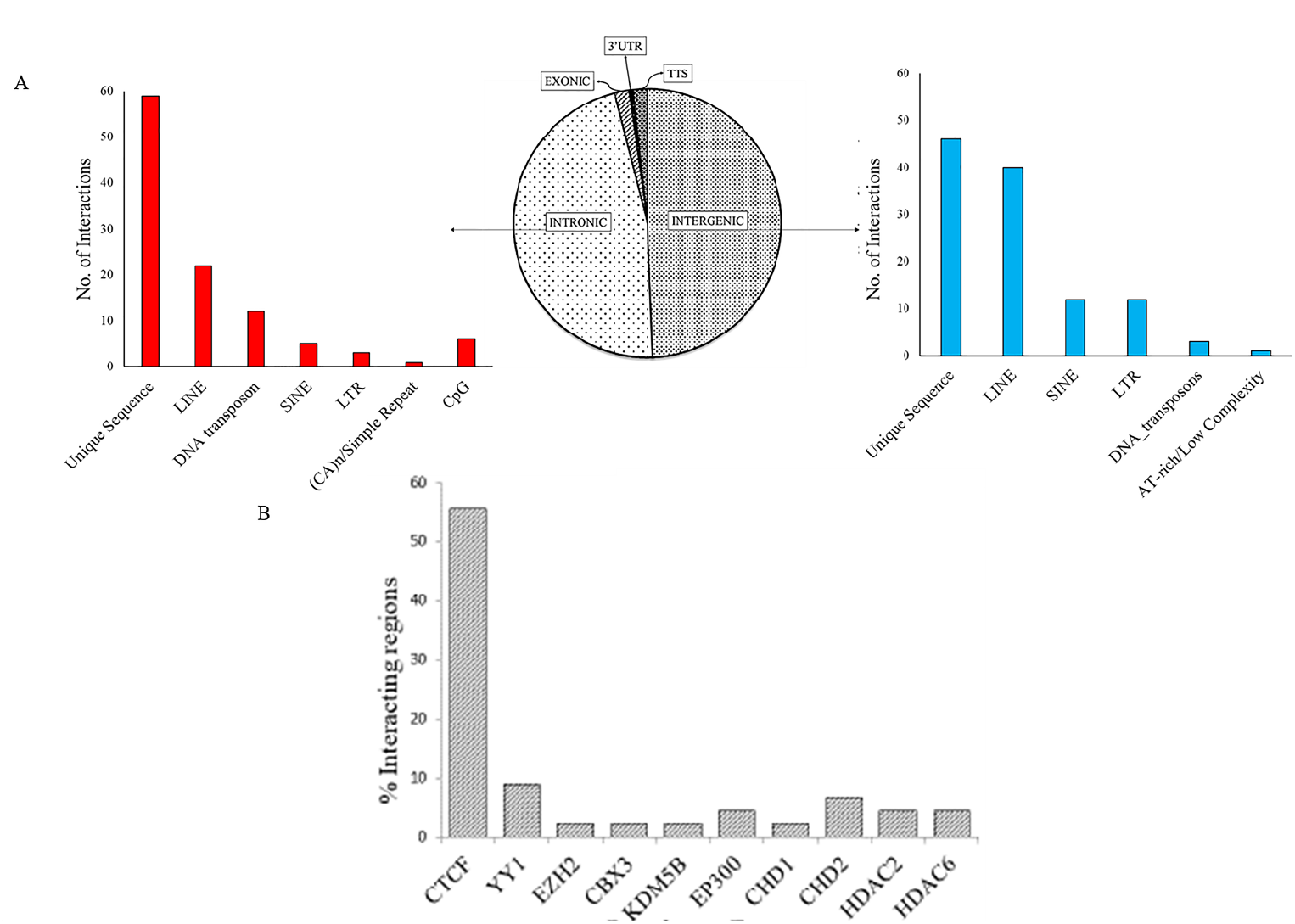
Association of the interactor sequences with cis-elements and repetitive sequences. A-The pie-chart indicates the distribution of interactor sequences within exonic, intronic and intergenic regions. The bar graphsshow the distribution of intronic sequences(red) and intergenic (blue)sequences into different classes of repetitive elements. B-Distribution (%) of interactor sequences on the basis of their enrichment with epigenetics factors (derived from ENCODE ChIP-Seq).

### 1.3.9 Identification of PcG-enriched interactors

Apart from the interactors identified by 4C-sequence, a number of potential intra-chromosomal interactors were identified on the basis of their enrichment with YY1, CTCF and other polycomb proteins, such as RBBP5 & PHF8 in ChIP-seq data from ENCODE (Figure 9B and 10A). These potential regulatory elements varied in their distance from the bait (PRE-PIK3C2B sequence), i.e. approximately 60 Kb to 3.8 Mb. The idea was to capture those interactions that are not easily identified in the 4C-sequencing analysis due to stringency criteria imposed. The sequences were identified, and labelled from 1 to 9 as shown in Figure 10A. Using 3C, we not only validated these interactions but also predicted the preferred topologies in which these interactions could occur (Figure 10C and Supplementary Figure S4). Two sets of primers are selected for each region, one set mapping to the bait and another mapping to the potential interactor sequence (in this case sequence 1). Following 3C, we validate the interactions by PCR using different combinations of primers. The primer-set combination where the amplicon is detected, is then used to map topologies. In case of sequence 1, two probable topologies exist as depicted in Figure 10C, based on the amplicons obtained using different primer sets. Using a similar approach, we validated and identified preferred/probable topologies, for regions 2,3 & 4 (Figure 11 & 12) and regions 5, 6, 7 & 8 (Supplementary Figure S5 & S6). The absence of amplicon in PCRs using control samples (non-ligated and gDNA), further confirms that these interactions are specific (Supplementary Figure S7).

**Figure 10.**
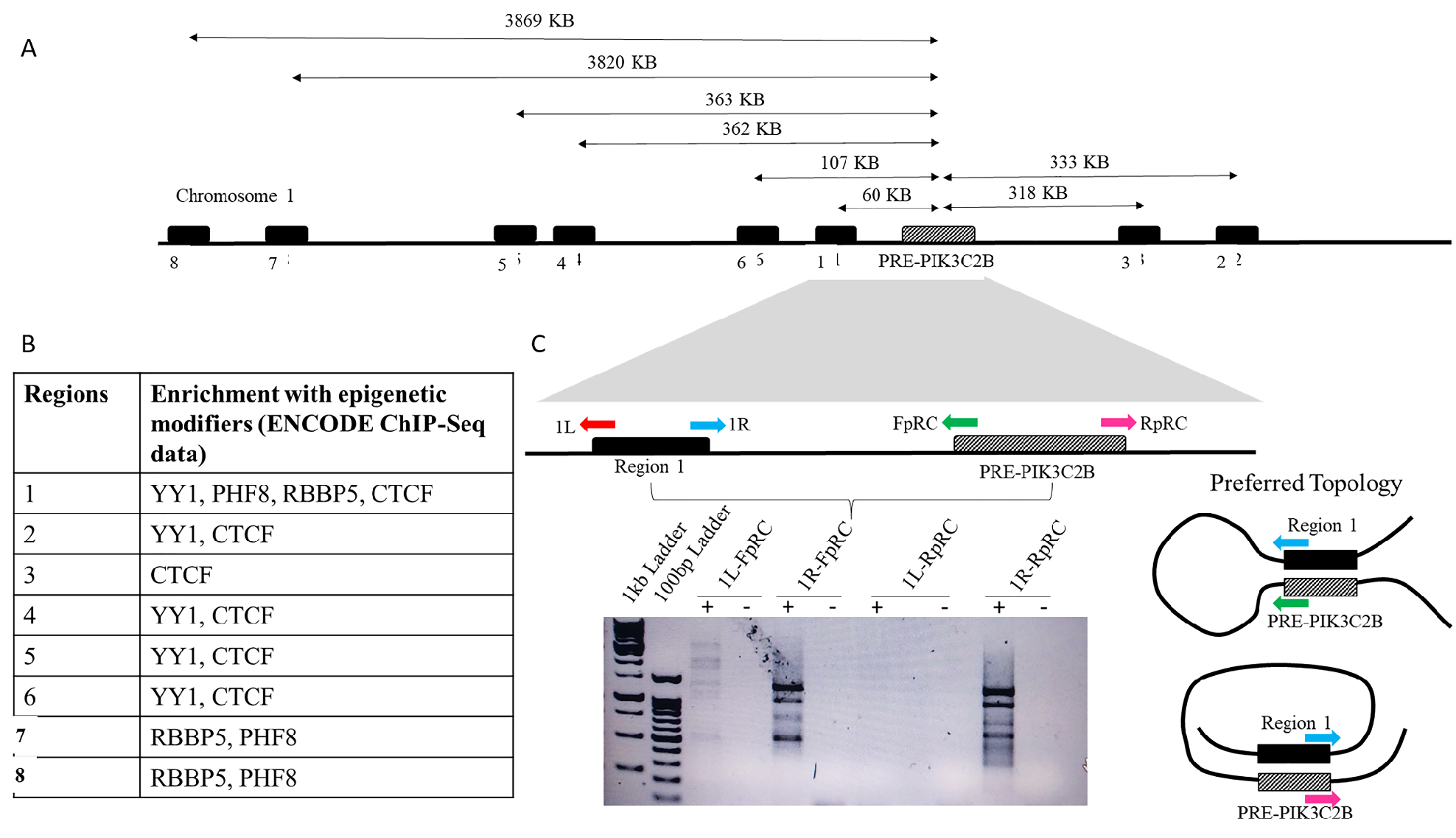
3C-PCR based *de-novo* identification of *PRE-PIK3C2B* associated chromatin interactions. A-region on the chr.1 with sites **(black box, 1-8**enriched with YY1, CTCF and other chromatin proteins. The distance (kb) of the sites from *PRE-PIK3CB* (striped box) are indicated (lines with double arrow). B-Table shows the ENCODE Chip-Seq based chromatin protein enrichment analysis, C-The line diagram showing the distance between *PRE-PIK3C2B* and site1. 1L,1R are the primers mapping to site 1 and FpRC, RpRC primers map to PRE-PIK3C2B. D-profile of the amplicons obtained with different primer sets. The gel image shows the amplicons indicating potential interactions.E-Topologies inferred from the amplicons obtained. (+)- indicates presence of the template, (-) – indicates the absence of the template.

**Figure 11.**
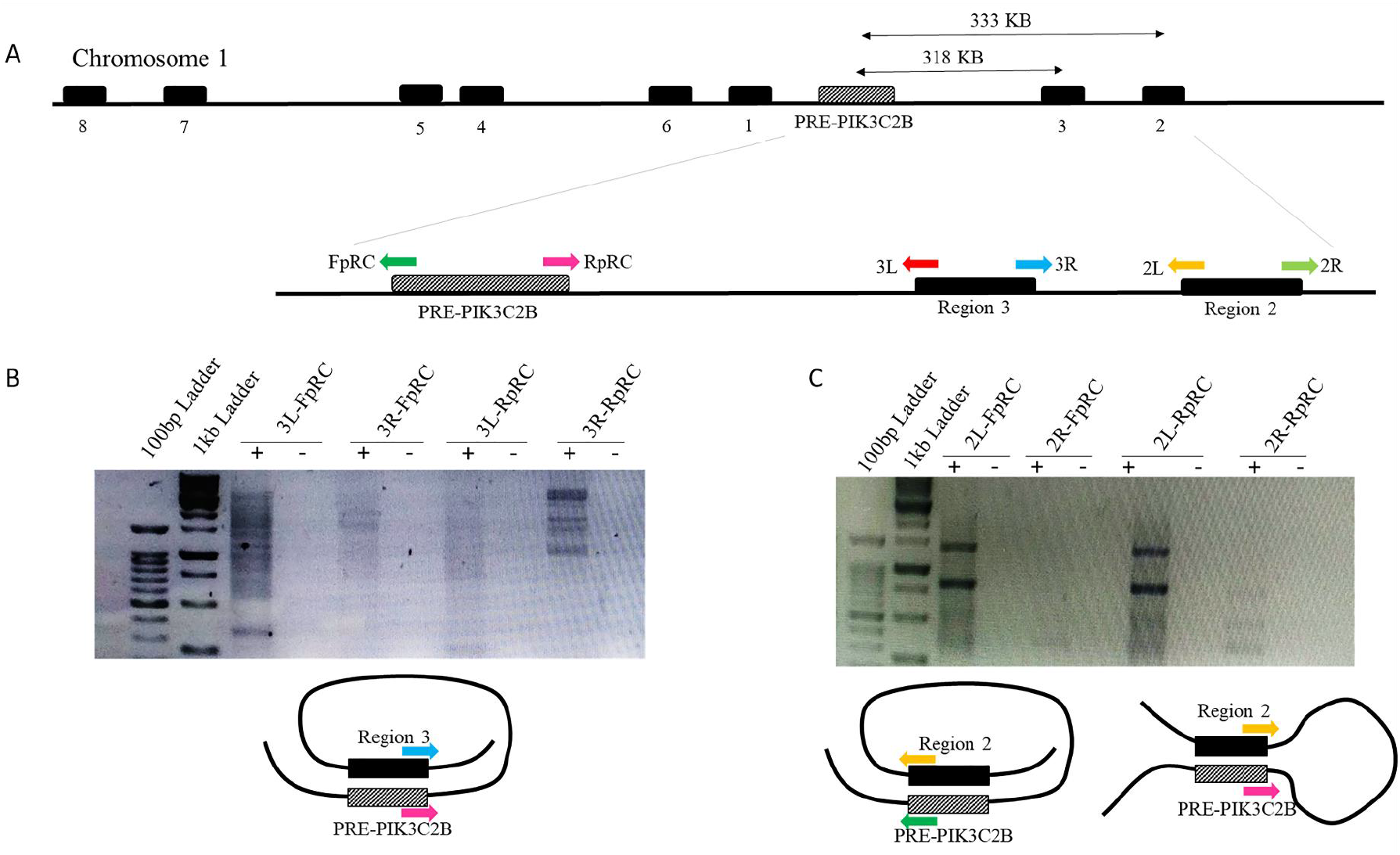
3C-PCR based *de-novo* identification of *PRE-PIK3C2B* associated chromatin interactions. A) Depicts a region on the chr1 with sites (black box) enriched with YY1, CTCF and other chromatin proteins. The double arrows indicate the distance between PRE-PIK3C2B and sites 2 and 3. The 2nd line diagram shows the distance between *PRE-PIK3C2B* and sits 2&3 along the position of primers used for 3C-PCR based identification of chromatin interactions. 2L/2R & 3L/3R are the primer sets mapping to site 2 and 3, respectively and FpRC/RpRC primers map to PRE-PIK3C2B. B) The Gel pic shows the amplicons indicating potential interaction between PRE-PIK3C2B and Site3. Based on the primer combination that gave the amplicon product, topologies of interaction between PRE-PIK3C2B and Site3 can be inferred. C) The electrophoretic profile image shows the amplicons indicating potential interaction between PRE-PIK3C2B and Site2. Based on the primer combination that gave the amplicon product, topologies of interaction between PRE-PIK3C2B and Site2 can be inferred. (+)- indicates presence of the template, (-) – indicates the absence of the template.

**Figure 12.**
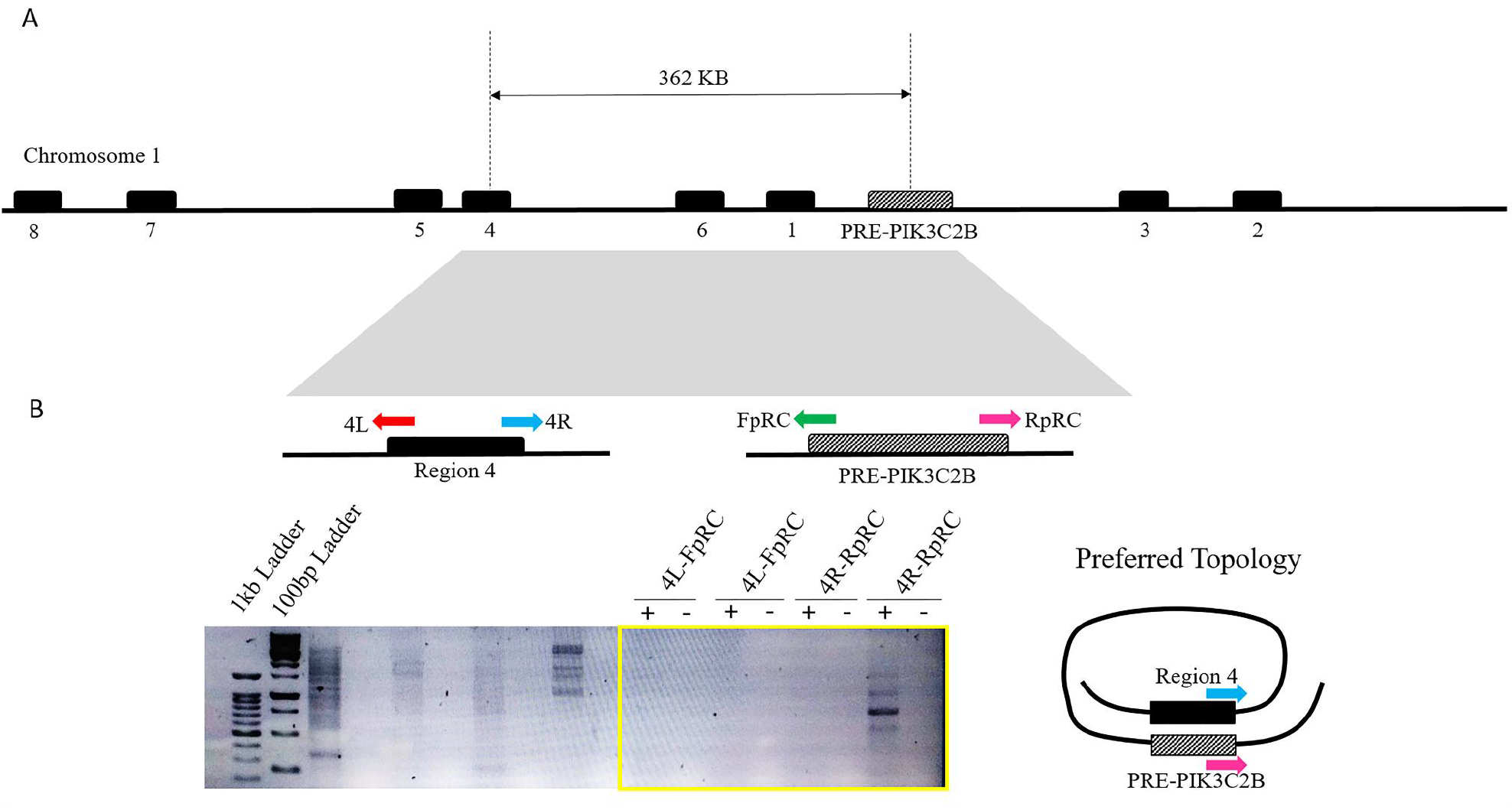
3C-PCR based *de-novo* identification of *PRE-PIK3C2B* associated chromatin interactions. A) Depicts a region on the chr1 with sites (black box) enriched with YY1, CTCF and other chromatin proteins. The double arrow indicates the distance between PRE-PIK3C2B and site 4. The 2nd line diagram shows the distance between *PRE-PIK3C2B* and site 4 along the position of primers used for 3C-PCR based identification of chromatin interactions. 4L/4R is the primer set mapping to site 4 and FpRC/RpRC primers map to PRE-PIK3C2B. B) The Gel image shows the amplicons indicating potential interaction between PRE-PIK3C2B and Site4. Based on the primer combination that gave the amplicon product, topologies of interaction between PRE-PIK3C2B and Site4 can be inferred. (+)- indicates presence of the template, (-) – indicates the absence of the template.

## 1.4 Discussion

The formation of the higher-order chromatin structure and its 3D architecture plays a crucial role in regulating transcription. These topologies result in the formation of DNA loops bringing the target genes closer to the distal enhancers, silencers and insulators [19]. The PRE mediated looping for gene silencing bringing the polycomb response element closer to the promoter and 5’-3’ gene looping for transcription termination are known [19]. The promoter-enhancer interaction at the beta-globin locus is one of the early examples of long-range interaction in human cells [37]. Further, it is reported that a group of genes can co-occupy sites that are foci of Polycomb proteins in *Drosophila* [38].

Previous study from our lab demonstrated the coordinated expression of *PIK3C2B* and its flanking genes *MDM4* and *RRR1R15B* in U87 and HEK cells lines [30]. *MDM4* is 21.4 kb and *RRR1R15B* is 11 kb from *PRE-PIK3C2B*. PRE-PIK3C2B, apart from interacting with polycomb proteins, also functions as a TRE without any sequence alteration, and the three genes in terms of their expression pattern, respond similarly to the relative concentration of YY1, MLL and Th-POK [30]. In this current study, we demonstrate the long-range interaction of *PRE-PIK3C2B*, not only with intra-chromosomal regions but also inter-chromosomal sites. To our knowledge, this is one of the early studies of chromatin interactions associated with a PRE in Humans. We find the predicted loops formed by the interactors of PRE-PIK3C2B encompass TAD (Topologically Associated Domains) sequences. TADs, divide the genome into physical domains and were detected by high resolution chromatin interaction maps in metazoans. Within the TADs the genomic interactions are strong and large in numbers. But there is a sharp decline in the number and strength of genomic interactions between TADs [39]; [40]; [41].

The formation of chromatin loops depends on proteins such as CTCF, Cohesins and the Mediator Complex. The detection of binding sequences for these factors in the regions of interaction with PIK3C2B suggests its involvement in chromatin domains. The mediator complex mainly interacts with enhancer-promoter and promoter-promoter loops and play an important role in gene transcription whereas CTCF participates in longer-range interactions [42], [43]; [44]. CTCF is generally associated with the TAD boundaries and thus, maintain overall architecture of TADs. The cohesins, present a ring structure, to bind the chromatin loops and are known to associate with both the mediator complex and CTCF [45]; [44].

The number of inter-chromosomal interactors (~209) identified was larger than that of intra-chromosomal interactors (~22). A large number of interactor sequences mapped on the chromosome 8 (24 interactions) and the least on Chr19 and 22 (2 interactions), with no interactions on ChrY. Chr8 has a gene density of 5.6 gene/Mb which is less than the average (~10 gene/Mb). Nearly half of chromosome 8 is composed of repetitive sequences with 44.5% comprising of transposable elements [46]. 8% of genes on Chr8 participate in brain development. Chr19, in contrast, has highest gene density of all chromosomes (26 genes/Mb) which is double the genome-wide average [47]. Although, Chr19 also has high density of repeats as well as unusually high G+C content (48%) **[47].** Our observation establishes a direct correlation between repetitive sequences and the number of long-range interactions. This observation is further supported by our analysis where a large proportion of the interactor sequences are derived from LINE and SINE elements.

A number of intra-chromosomal interactions might be false positive due to the close proximity of the regions to the bait (often called *cis* interactions), although cis-distal interactions are known [48].

Therefore, in order to identify other interacting regions, we applied a criterion for enrichment of polycomb complex proteins like YY1, EZH2, SUZ 12 and CTCF on the regions proximal to *PRE-PIK3C2B*, using ChIP-Seq data from ENCODE data. These regions were not identified in 4C analysis. This led us to detect and validate eight sites, as interactors. It is possible that these sites play a role in coordinated expression of *PIK3C2B, MDM4* and *PPP1R15B*.

It is well-known that various genes and regulatory elements are brought together to form repressed or active chromatin hubs to facilitate the coordinated gene activity [49]; [50]. There are a number of examples in *Drosophila*, of PREs participating in 3D chromatin organization in order to maintain transcription states. *Fab-7* PRE, located between the Abd-A and abd-B transcription units, when present in multiple copies, initiates long range gene contacts that play an important role in PcG mediated silencing [51] . The genes of the Antennapedia and Bithorax complexes in *Drosophila* interact within the Polycomb bodies and these interactions were shown to be tissue specific. [52] reported EZH2 dependent long-range interaction and demonstrated that these interactions are conserved in *Drosophila virilis*, which is a distantly related species [38]. Our results strongly support such interactions in the human genome and we detected the enrichment of polycomb complex members around this region in ENCODE data. Almost 58% (134/231) of the interactors mapped within +/- 2Mb of the TSS, suggesting the potential influence of the long-range interaction of PRE-PIK3C2B, on transcription. We observed that there is uniform low expression of these genes. The gene expression meta-analysis clearly indicates that the genes associated with interactor sequences are co-expressed or co-repressed, this is in comparison to the uncoordinated expression observed for the random set of genes used as control for this analysis.

A significant proportion of the DNA interactors map to repetitive regions and a number of interactors in genic regions map within the LINE elements. This is not surprising since there are examples of a number of genes harboring repetitive elements. Our analysis has shown that *hPRE-PIK3C2B* itself is derived from LINE-1 elements (unpublished results). We find an over-representation of LINE and SINE elements within the repetitive sequences in the interactors. The LINE elements are implicated in heterochromatin organization. Initially repetitive elements were known to be enriched mainly with PcG-mediated H3K27me3 mark, as a compensatory silencing mechanism. However, the suppression of mobilization of the endogenous murine leukemia virus elements by polycomb complex and the demonstration of PRC2 and PRC1 complexes targeting the genomic repeats indicate the role of polycomb complexes in silencing mechanisms [53].

The interactors we identified, are enriched with SINE elements. YY1, a member of the Polycomb Repressive Complex 2, is known to bind to Alu-consensus motif (CCATGTTGGC), which represents >30% of all Alu repeats (Oei et al., 2004). Repeats like LINE-1 and Alu participate in heterochromatin formation during X-chromosome inactivation [54]. Interestingly the length of the repeat is correlated with the expression status of the associated gene. The DXZ4 and D4Z4 macro-satellites map to the X chromosome and show alternative epigenetic states that depend on the number of repeats. Unlike PRE-PIK3C2B where the PRE and TRE like activity is correlated with relative abundance of the protein factors, the activation function of D4Z4 is correlated with the reduction in the copy number of D4Z4 [55], [31]. As a consequence of this, genes in 4q35 region are activated in patients of FSHD (Facioscapulohumeral Muscular Dystrophy), raising the possibility of long-range interaction leading to altered transcription profile.

In Glioblastoma, 1q32 region harboring the *PIK3C2B* gene, is amplified [56]; [57] and the expression of *PIK3C2B* plays a major role in resistance to erlotinib, an anti-cancer drug. However, the correlation between amplification of 1q32 and increased expression of *PIK3C2B* is not clear. Thus, it would be interesting to know, how the multiple copies *PRE-PIK3C2B* in 1q32 amplification along with the associated chromatin interaction regulate *PIK3C2B* expression.

Our analysis based on ENCODE ChIP-Seq, shows that in K562 and HepG2 cell lines, the interactor sequences (and their flanking regions) are enriched with CTCF and YY1 proteins. The interactor sequences in K562 cell line are enriched with YY1, CTCF and CBX3 along with H3K27me3 and this establishes a strong correlation between histone modification and enrichment of epigenetic modifier. A number of PREs harbor YY1 binding sites and majority of the *Drosophila* PREs are identified on the basis of density of Pho (YY1) binding sites. The Human PRE-PIK3C2B contains a 25mer (with YY1 binding site) repeated 25 times and binds to YY1 [29], [30]. The mouse PRE-kr and D11.12 element, in the human HOXD cluster contain YY1 binding sites [58]; [59]. By mutating the minimal binding site for YY1 (GCCAT), it was shown that BMI1 enrichment was impaired even though SUZ12 binding was unaffected [59]. Thus the presence of YY1 is a strong marker for identification of PRE sequences.

In contrast, the interactor sequences in HepG2 cell lines show an enrichment of H3K4me3, as active histone mark, along with strong YY1 and CTCF enrichment. There are a number of studies that have shown the activating function of YY1. A recent study has shown that YY1 maintains pluripotency by activating gene expression, by interacting with BAF complex and also the enrichment of H3K4me3 [60]. In another study, YY1 was shown to preferentially bind to Trithorax Response Elements and not Polycomb Response Elements thus acting as potential recruiter of MLL1/2-TrxG Complex [61].

This was in concurrence with the report of [62], where the interaction of YY1 with active promoters, but not PcG target genes was demonstrated. In primate genome, CTCF-YY1 co-occupy regions, that are known to be promoter-proximal and gene-distal and are enriched for RNA polymerase II, transcription factor binding and active histone marks [63]. In a recent report, transposable elements (TE) were established as important contributors to divergent chromatin looping in mouse and humans [64]. According to this study, ~⅓ of the CTCF-binding sites are contributed by the TE activity in humans and mouse genome [64]. In the light of these observations, the coordinated expression of *PIK3C2B* and its neighboring genes *MDM4* and *PPP1R15B* as well as the genes associated with the interactor sequences (inter-chromosomal interactions) may be mediated by the TE enriched regions along with the CTCF-binding sites.

In summary our 4C analysis has led to the identification of the long-range interactions of a human PRE. It would be interesting to examine if *hPRE-PIK3C2B* along with these domains form a part of the repression hub. Further validation and analysis is required to decipher if the long range interaction of PRE as seen in the case of *hPRE-PIK3C2B*, is important for architectural integrity of the nucleus as well as the coordinated, extended regulatory end-effects.

## Acknowledgements

VB gratefully acknowledges the financial support from SAP-II from University Grants Commission (UGC) and SERB No.60(0102)/12/EMR-II for financial support through research grant. JM acknowledges finanacial support from SERB No.60(0102)/12/EMR-II through Research Associate fellowship. We acknowledge the Bioinformatics facility at ACBR supported by the Department of Biotechnology, Government of India.

## Supplementary Figure Legends

**Supplementary Figure 1.**
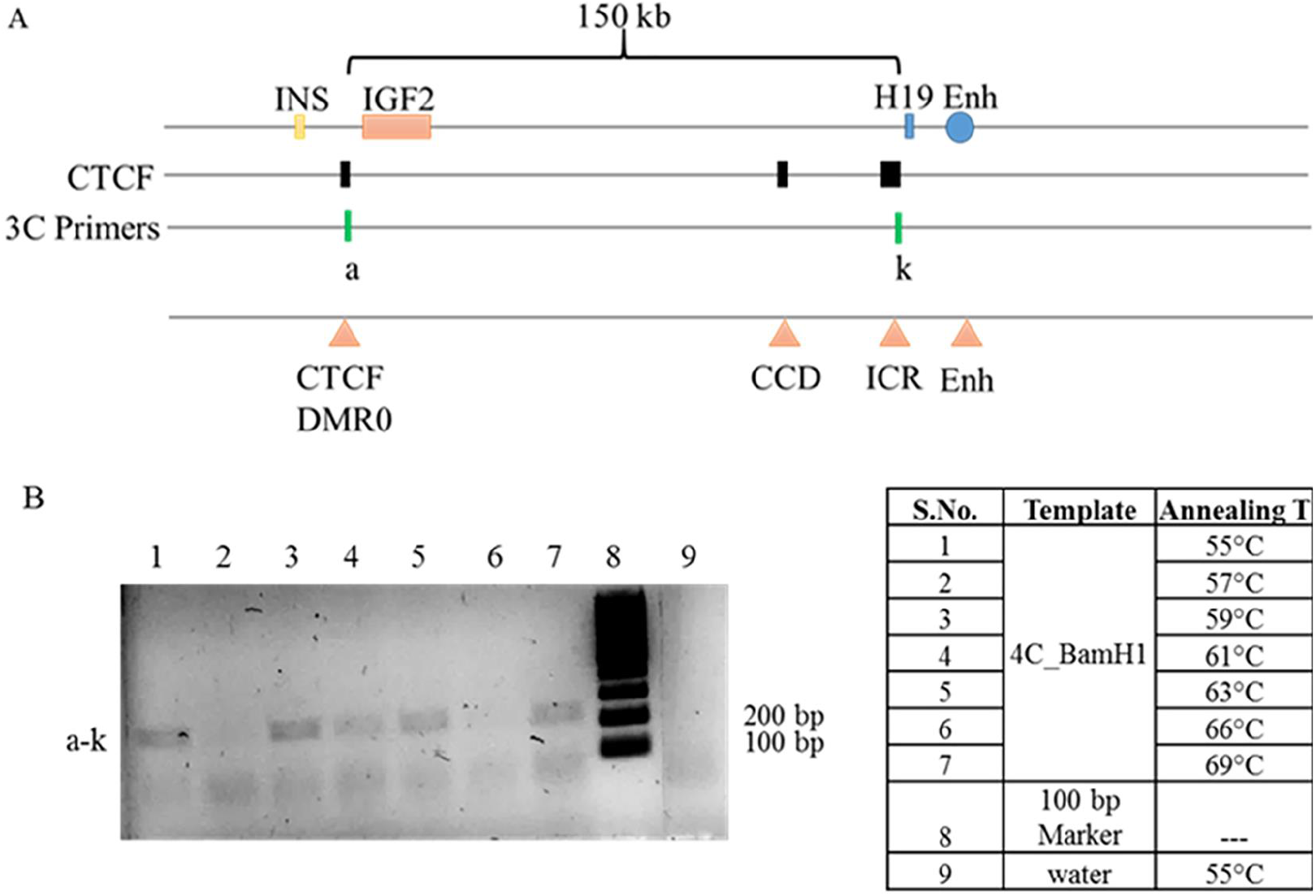
Validation of 4C-protocol using known interactions at the IGF2-H19 locus. (A) The 3C primers (a and k) mapping to differentially methylated regions near IGF2 and H19 loci enriched with CTCF binding were used as positive controls. (B) The bands indicate the interactions detected in the 4C experiments performed with BamH1 digestion followed by ligation. Different annealing temperatures were used to standardize the PCR reactions. The negative PCR control was clean (lane 9).

**Supplementary Figure 2.**
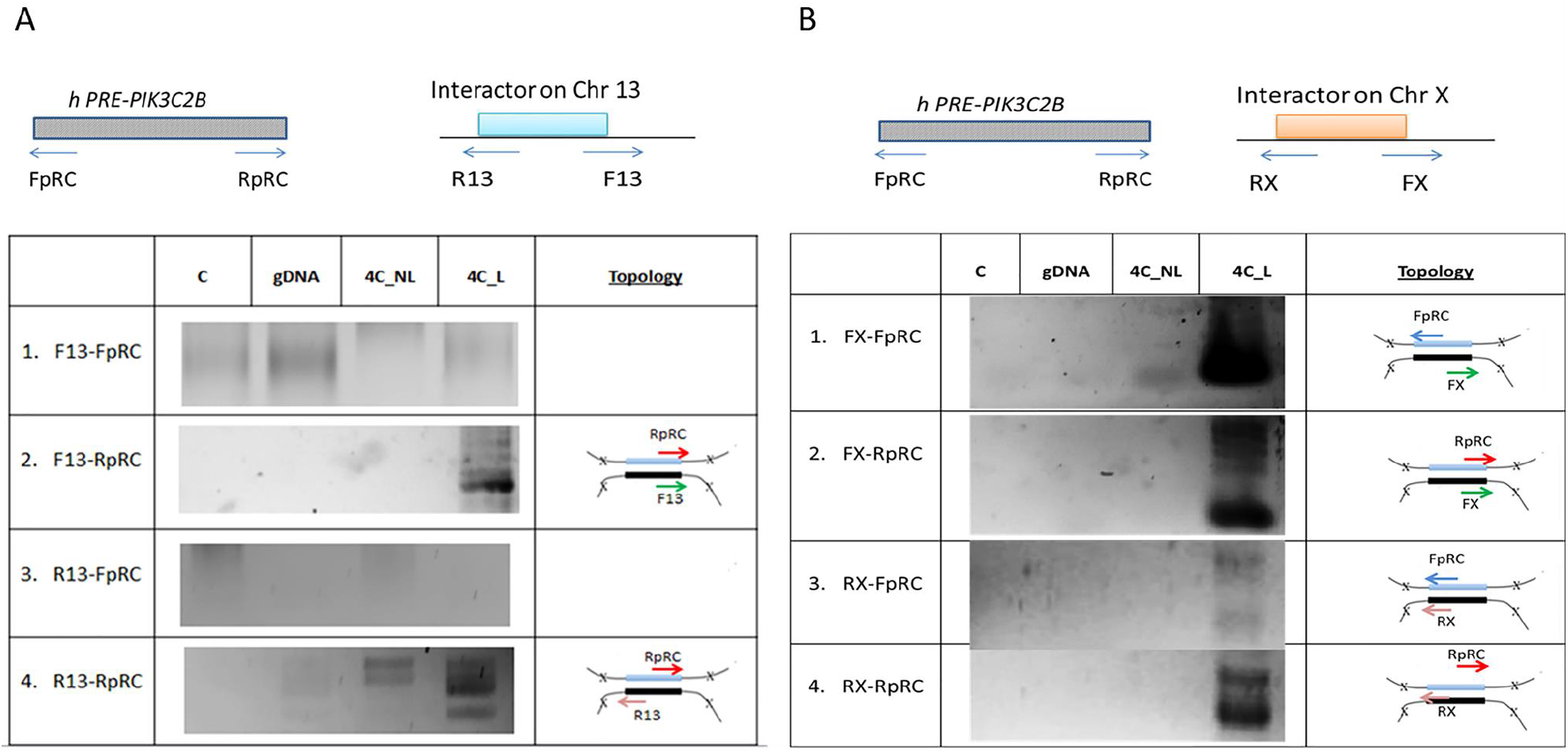
3C-PCR based validation of the inter-chromosomal interactions identified in 4C-Seq. The interactor sequences are located on Chr13 and X. A) The tabular depiction indicating the primer combinations used for validation of interaction between PRE-PIK3C2B and site on chr13 along with the gel image (C-PCR negative control, 4C_L-ligated, 4C_NL-non-ligated, gDNA-demonic DNA restriction digested and ligated) and preferred topologies. B) The tabular depiction indicating the primer combinations used for validation of interaction between PRE-PIK3C2B and site on chrX along with the gel-picture (C-PCR negative control, 4C_L-ligated, 4C_NL-non-ligated, gDNA-demonic DNA restriction digested and ligated) and preferred topologies.

**Supplementary Figure 3.**
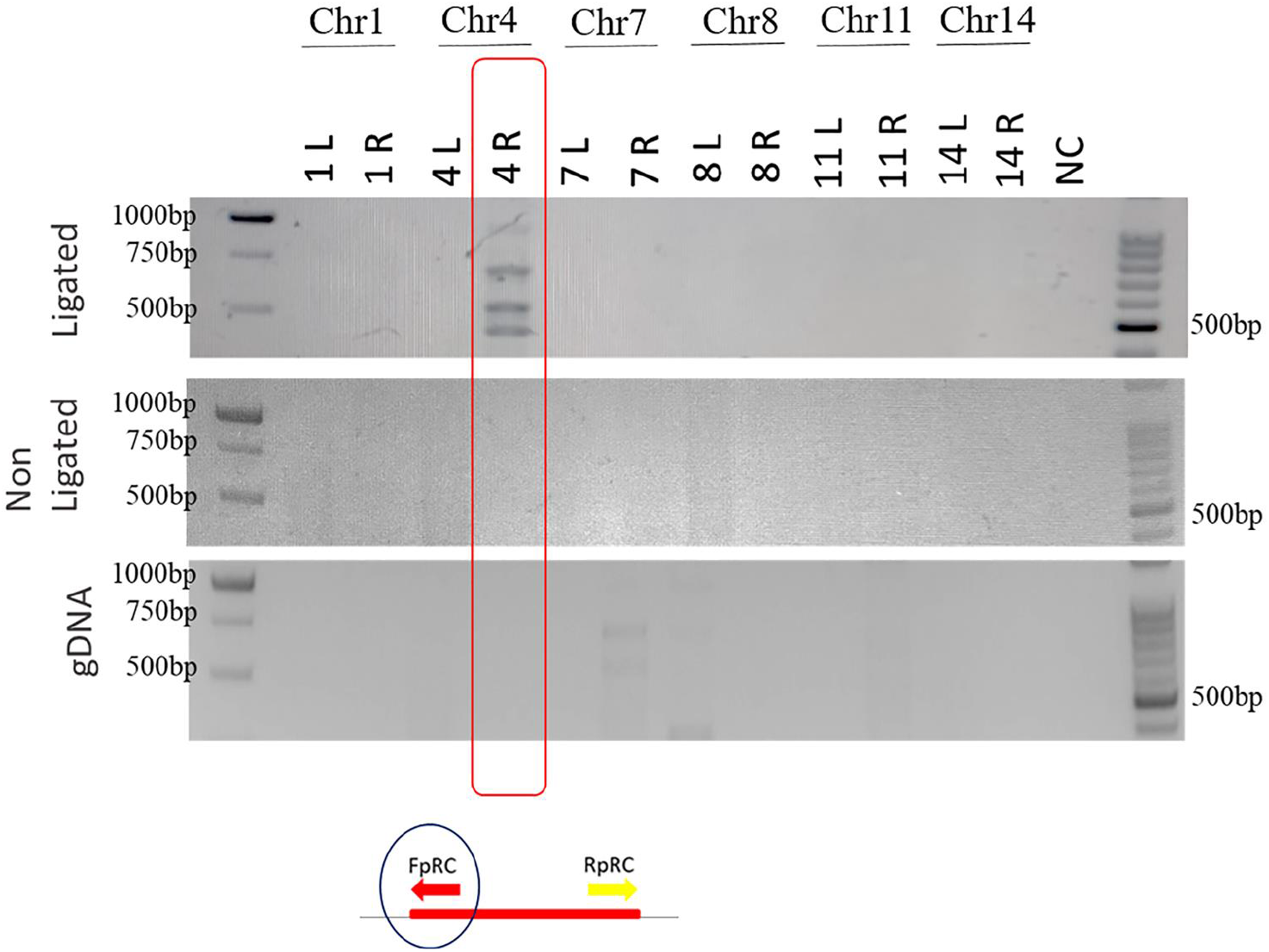
3C-PCR based validation of intra- and inter-chromosomal interactions (chr 1, 4, 7, 8, 11 and 14) identified in 4C-Seq. The top gel picture shows PCR product amplified using the 4C-ligated sample. The middle gel picture shows the PCR amplification of products using 4C non-ligated control samples. The lower gel picture shows the PCR amplification of products using 4C gDNA control samples.

**Supplementary Figure 4.**
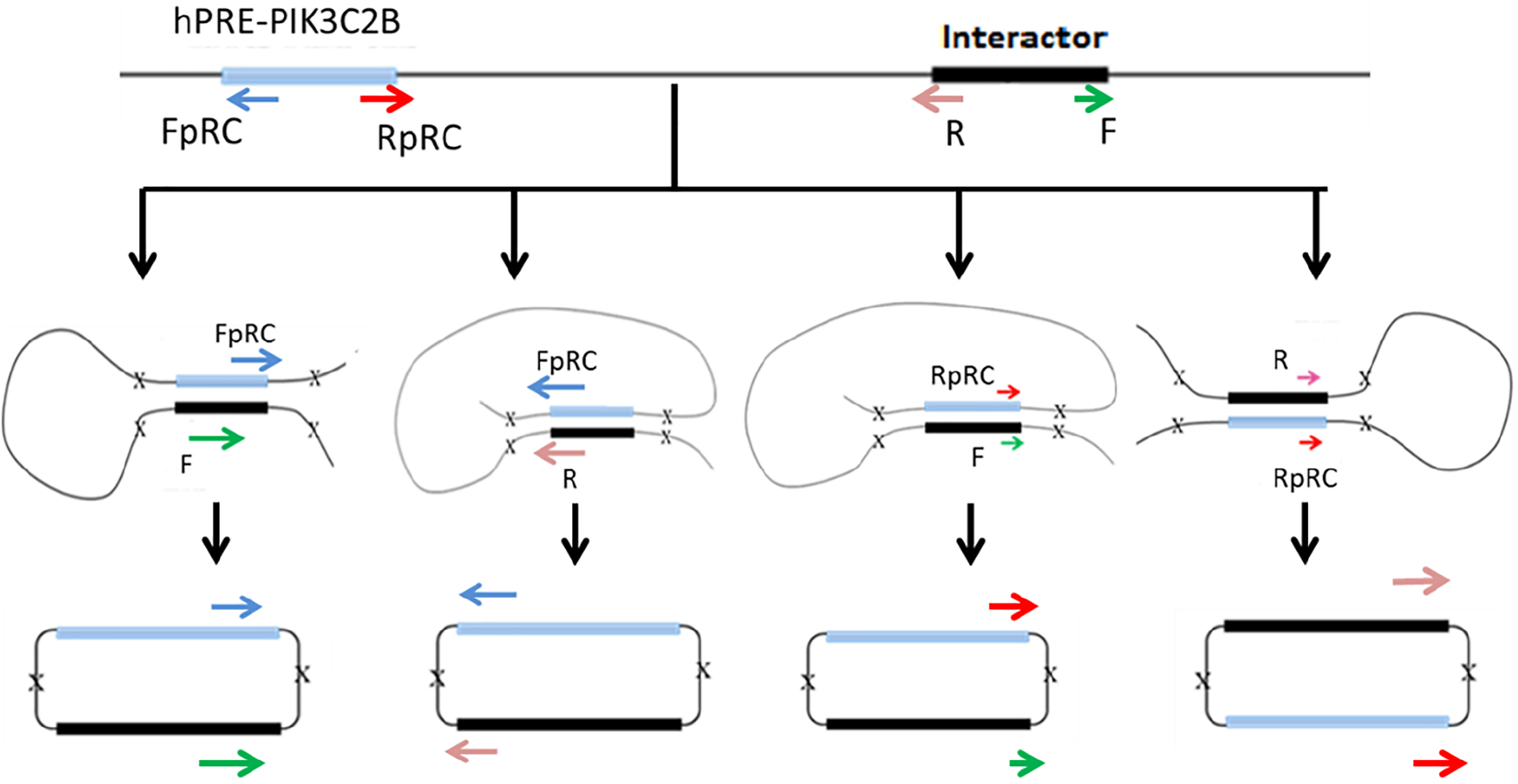
Schematic diagram howing the positioning of the primers for validation using 3C approach. The primer mapping to the query interactors (pink and green arrowheads) were designed to account for the 4 potential ways by which the query sequences and the bait that is the *PRE-PIK3C2B* come together. The primers FpRC/RpRC mapping to the bait [PRE-PIK3C2B (blue and red arrowheads)] were the same as the one used for 4C sequencing.

**Supplementary Figure 5.**
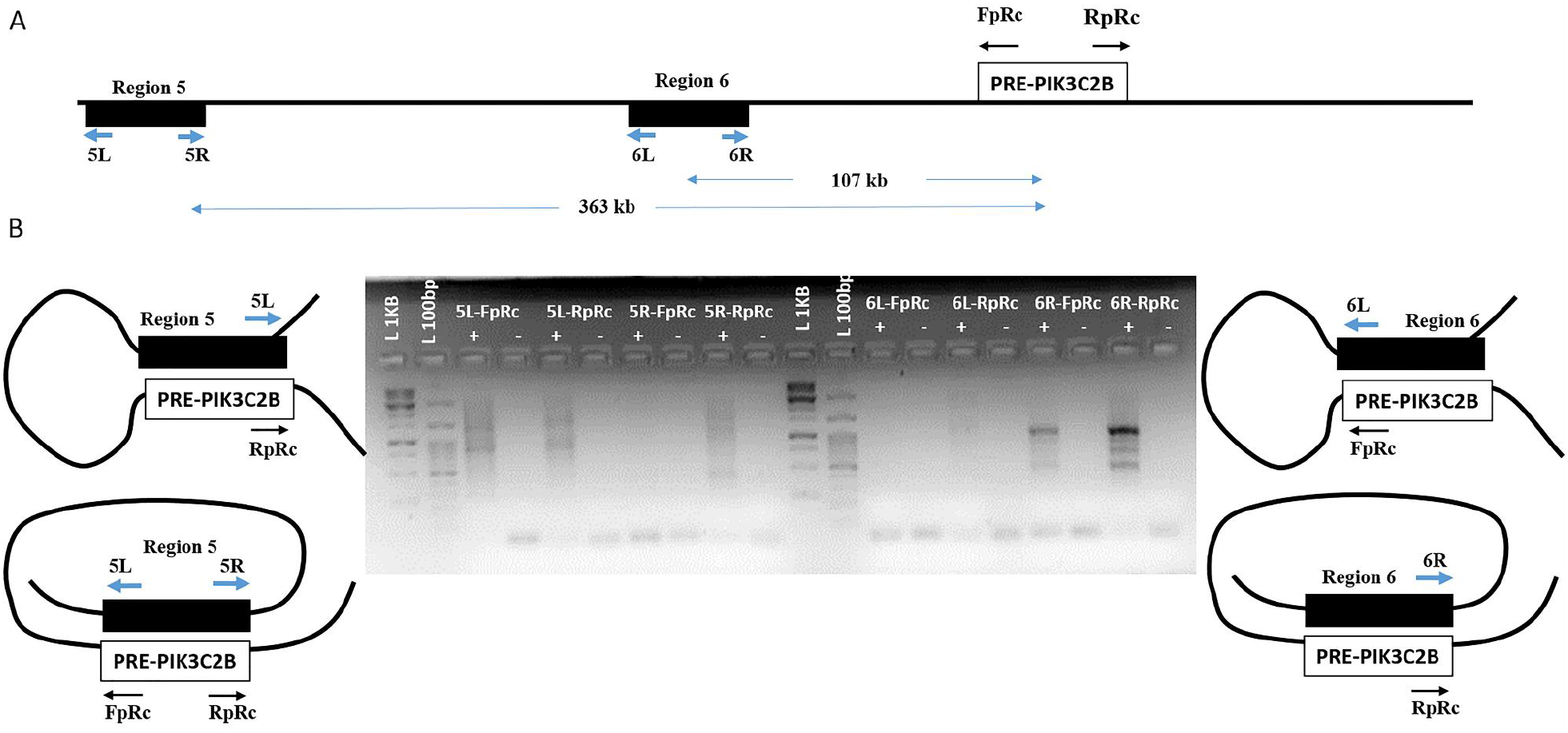
3C-PCR based *de-novo* identification of *PRE-PIK3C2B* associated chromatin interactions. A) The line diagram depicts regions 6 and 5 on the chr1 with sites (black box) enriched with YY1, CTCF and other chromatin proteins. The double arrows indicate the distance between PRE-PIK3C2B and sites 5 and 6. 5L/5R & 6L/6R are the primer sets mapping to site 5 and 6, respectively and FpRC/RpRC primers map to PRE-PIK3C2B. B) The Gel image shows the amplicons indicating potential interaction between PRE-PIK3C2B and sites 5 & 6. Based on the primer combinations that gave the amplicon product, topologies of interaction between PRE-PIK3C2B and sites 5&6 can be inferred.

**Supplementary Figure 6.**
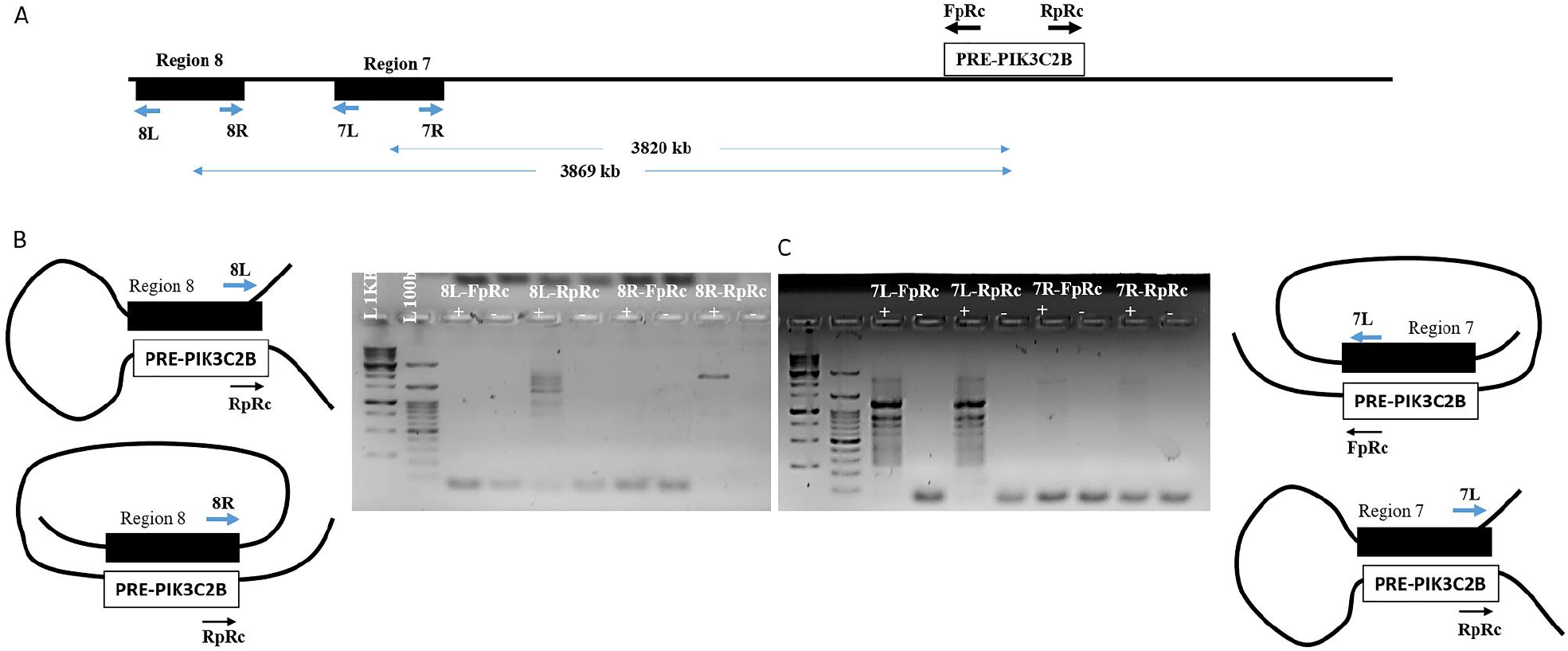
3C-PCR based *de-novo* identification of *PRE-PIK3C2B* associated chromatin interactions. A) The line diagram depicts regions 7 and 8 on the chr1 with sites (black box) enriched with YY1, CTCF and other chromatin proteins. The double arrows indicate the distance between PRE-PIK3C2B and sites 7 and 8. 7L/7R & 8L/8R are the primer sets mapping to sites 7 and 8, respectively and FpRC/RpRC primers map to PRE-PIK3C2B. B) The Gel image shows the amplicons indicating potential interaction between PRE-PIK3C2B and sites 7 & 8. Based on the primer combinations that gave the amplicon product, topologies of interaction between PRE-PIK3C2B and sites 7 & 8can be inferred.

**Supplementary Figure 7.**
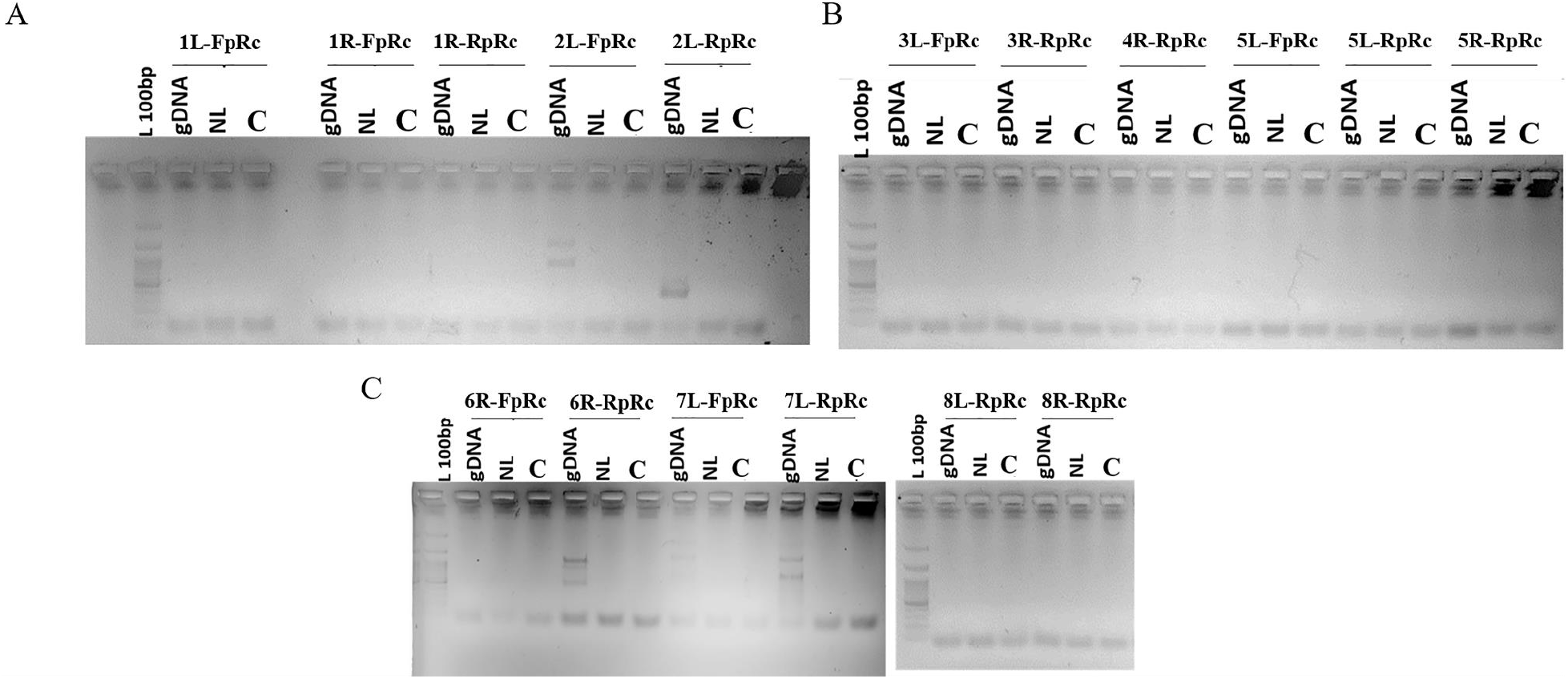
Control PCRs for 3C-PCR-based de-novo identification of PRE-PIK3C2B associated chromatin interactions. A) Gel picture for the 3C-PCR using non-ligated (NL) and gDNA controls as templates for identifying chromatin interactions between chr1 & 2 and PRE-PIK3C2B. −ve – negative control for PCRs (with no template). B) Gel picture for the 3C-PCR using non-ligated and gDNA controls as templates for identifying chromatin interactions between chr3, 4 & 5 and PRE-PIK3C2B. −ve – negative control for PCRs with no template. C) Gel picture for the 3C-PCR using non-ligated and gDNA controls as templates for identifying chromatin interactions between chr6, 8 & 9 and PRE-PIK3C2B. C – negative control for PCRs with no template.

## Supplementary Tables

**Supplementary Table 1.**
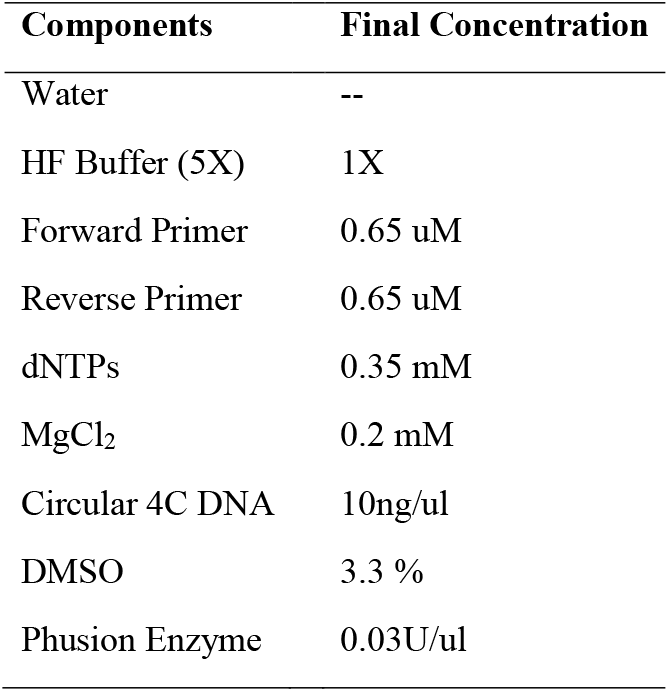
The following reaction was set up in total volume of 20ul:

| Components | Final Concentration |
| --- | --- |
| Water | -- |
| HF Buffer (5X) | 1X |
| Forward Primer | 0.65 uM |
| Reverse Primer | 0.65 uM |
| dNTPs | 0.35 mM |
| MgCl<sub>2</sub> | 0.2 mM |
| Circular 4C DNA | 10ng/ul |
| DMSO | 3.3 % |
| Phusion Enzyme | 0.03U/ul |

**Supplementary Table 2.**
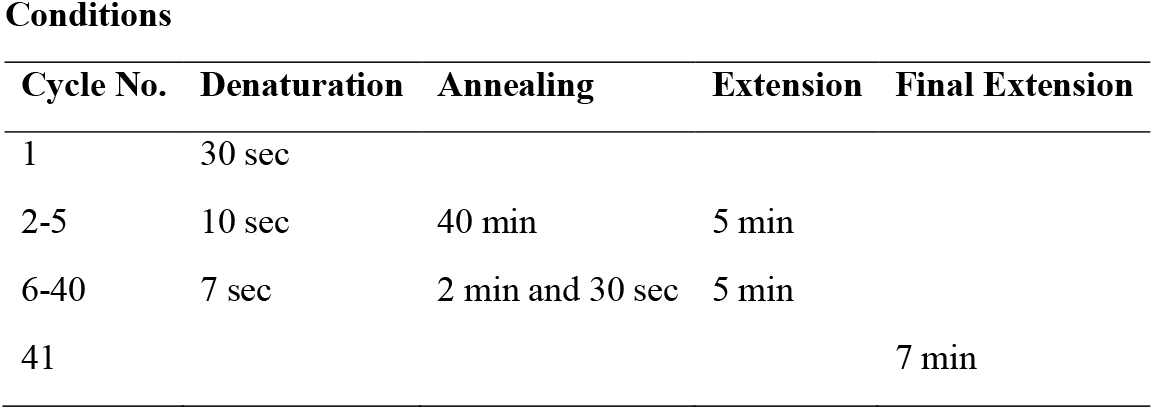

Conditions
| Cycle No. | Denaturation | Annealing | Extension | Final Extension |
| --- | --- | --- | --- | --- |
| 1 | 30 sec |  |  |  |
| 2-5 | 10 sec | 40 min | 5 min |  |
| 6-40 | 7 sec | 2 min and 30 sec | 5 min |  |
| 41 |  |  |  | 7 min |

**Supplementary Table 3.** For the 2^nd^ set of PCR, the following reaction was set up:

| Components | Amount per reaction (ul) |
| --- | --- |
| Water | 10.9 |
| HF/GC Buffer (5X) | 4.0 |
| 5 pmoles/ul Forward Primer | 1.3 |
| 5pmoles/ul Reverse Primer | 1.3 |
| 10mM dNTPs | 0.7 |
| MgCl<sub>2</sub> | 0.3 |
| 1:10 diluted PCR | 1.5 |
| Phusion Enzyme | 1.0 |

**Supplementary Table 4.**

Conditions (Running text)
| Cycle No. | Denaturation | Annealing | Extension | Final Extension |
| --- | --- | --- | --- | --- |
| 1 | 4 min |  |  |  |
| 2-34 | 30 sec | 1 min | 4 min |  |
| 35 |  |  |  | 7 min |

**Supplementary Table 5.** The Table list out the paired end fastq files sequenced on Illumina Miseq machine

| Files | Number of Reads |
| --- | --- |
| SO_5382_VBL_4C_gDNA_R1.fastq.gz | 553658 |
| SO_5382_VBL_4C_gDNA_R2.fastq.gz | 553658 |
| SO_5382_VBL_4C_L_R1.fastq.gz | 434284 |
| SO_5382_VBL_4C_L_R2.fastq.gz | 434284 |
| SO_5382_VBL_4C_NL_R1.fastq.gz | 542162 |
| SO_5382_VBL_4C_NL_R2.fastq.gz | 542162 |

**Supplementary Table 6.** The table list out the paired end fastq files sequenced on Illumina Nextseq machine

| Files | Number of Reads |
| --- | --- |
| SO_5382_VBL_4C_gDNA_R1.fastq.gz | 2316035 |
| SO_5382_VBL_4C_gDNA_R2.fastq.gz | 2316035 |
| SO_5382_VBL_4C_L_R1.fastq.gz | 2557052 |
| SO_5382_VBL_4C_L_R2.fastq.gz | 2557052 |
| SO_5382_VBL_4C_NL_R1.fastq.gz | 2637044 |
| SO_5382_VBL_4C_NL_R2.fastq.gz | 2637044 |

**Supplementary Table 7.** Combined Reads for different sample used in 4C sequencing analyses

| **Files** | **Number of Reads** |
| --- | --- |
| SO_5382_VBL_4C_gDNA_R1.fastq.gz | 2869693 |
| SO_5382_VBL_4C_gDNA_R2.fastq.gz | 2869693 |
| SO_5382_VBL_4C_L_R1.fastq.gz | 2991336 |
| SO_5382_VBL_4C_L_R2.fastq.gz | 2991336 |
| SO_5382_VBL_4C_NL_R1.fastq.gz | 3179206 |
| SO_5382_VBL_4C_NL_R2.fastq.gz | 3179206 |

